# Parietal–prefrontal feedforward connectivity in association with schizophrenia genetic risk and delusions

**DOI:** 10.1101/801506

**Authors:** Danielle Borrajo, Michelle La, Shefali Shah, Qiang Chen, Karen F Berman, Daniel R Weinberger, Hao Yang Tan

**Author notes:** Corresponding Author: Hao Yang Tan, Address: 855 North Wolfe Street, Suite 300, 3rd Floor, Baltimore, MD 21205.

## Abstract

**Background:** Conceptualizations of delusion formation implicate, in part, deficits at feed-forward information transfer across posterior to prefrontal cortices, resulting in dysfunctional integration of new information in favor of over-familiar prior beliefs. Here, we used functional MRI and machine learning models to examine feedforward parietal-prefrontal information transfer in schizophrenia patients in relation to delusional thinking, and polygenic risk for schizophrenia.

**Methods:** We studied 66 schizophrenia patients and 143 healthy controls as they performed context updating during working memory (WM). Dynamic causal models of effective connectivity were focused on prefrontal and parietal cortex, where we examined parietal-prefrontal connectivity in relation to delusions in patients. We further tested for an effect of polygenic risk for schizophrenia on connectivity in healthy individuals. We then leveraged support vector regression models to define optimal normalized target connectivity tailored for each patient, and tested the extent to which deviation from this target predicted individual variation in delusion severity.

**Results:** In schizophrenia patients, updating and manipulating context information was disproportionately less accurate than was WM maintenance, with a task accuracy-by-diagnosis interaction. Also, patients with delusions tended to have relatively reduced feedforward effective connectivity during context updating in WM manipulation. The same parietal-prefrontal feedforward prefrontal effective connectivity was adversely influenced by polygenic risk for schizophrenia in healthy subjects. Individual patients’ deviation from predicted ‘normal’ feedforward connectivity based on the support vector models correlated with delusional severity.

**Conclusions:** These computationally-derived observations support a role for feed-forward parietal-prefrontal information processing deficits in delusional psychopathology, and in genetic risk for schizophrenia.

## Introduction

Delusions occur across a range of psychotic disorders, and are characterized by distressing false beliefs that preoccupy, are held with conviction, and disrupt lives. Delusions have complex phenomenology, and possibly just as complex theories of causative and maintenance factors (1-5). Recent conceptualizations of the cognitive architecture of delusions have utilized Bayesian computational approaches to model how aberrant beliefs are over-learnt and become immutable (1, 6-8). It has been argued that the abnormal preoccupation and conviction with which beliefs are held may stem from cognitive biases against incorporating into beliefs of evidence otherwise (2, 4). These biases and erroneous beliefs are potentially further reinforced through exaggerated subcortical dopaminergic reinforcement (9, 10). Computationally, cognitive biases in the psychotic brain has been conceptualized as sub-optimal under-estimates of precision of likelihoods associated with new context information from the changing environment. Such under-estimation then leads to over-estimated precision and confidence about prior contexts or beliefs, and to resultant failures in arriving at inferences and behavioral adjustments to new (e.g. contrary) information and contexts, favoring instead, prior expectations and beliefs (6, 8). Consistent with this formulation, hallucinatory phenomena in early psychosis or at-risk mental states were recently observed to result from biasing prior expectations over incoming (contrary) sensory evidence (11). At the level of brain function, these computational biases appear to result from deficits in feedforward information processing, and have also been found in schizotypy and subclinical psychotic experiences (11). Related work on experimentally induced hallucinatory states has extended these findings to computational aspects of functional brain activation associated with hallucinations (12).

More specific to the abstract information processing in delusional beliefs, we previously found that Bayesian underweighting of new information in favor of prior beliefs was also associated with reduced feed-forward effective connectivity of information processing across cortical hierarchies from posterior cortical to prefrontal cortex (1). In patients with schizophrenia who had delusions, these findings were associated with subsequent over-reinforcement of dysfunctional prior belief models through subcortical connectivity with dopamine-rich loci in the midbrain (1). These changes were unlikely to be epiphenomena of treatment, because analogous information processing changes also were observed in unaffected siblings of patients with schizophrenia (1).

Our goal herein is to extend these recent observations linking delusional psychopathology, genetic risk for schizophrenia, and deficits in feedforward parietal-to-prefrontal connectivity during updating of new context information (1). Recent models of delusional and perceptual psychopathology suggest deficits in feedforward cortical integration of new information into belief models, leading to dysfunctional strong priors and consequent belief and perceptual disturbances (1, 12). Thus, we hypothesize that these effects should also be reflected in updating and manipulating context information in working memory (WM). We reason that because these WM processes involve updating prior information with new content, they should be critically engaged in the updating of new information into belief models putatively dysfunctional in delusions, and should also be associated with the underlying genetic mechanisms of schizophrenia (1, 12). We, therefore, predict that feedforward parietal-prefrontal connectivity while manipulating prior information held in WM might also relate to delusional psychopathology in patients with schizophrenia. Further, if these neural functions relate to genetic mechanisms of illness rather than confounds associated with medications or chronic ill health, we would expect that there would be relationships between feedforward parietal-prefrontal connectivity with polygenic risk for disease, particularly in healthy individuals.

We also aim herein to examine a substantial issue related to inter-individual variability in cortical function and connectivity, which may be particularly pronounced in the manner of prefrontal cortical processing we examine here (13). As each patient’s optimal neural function for a given cognitive task likely varies, a given patient’s deviation from his or her theoretical optimum should closely relate to cognitive dysfunction and psychopathology, if indeed, the specific neural function is implicated in these disease mechanisms. To examine this, we leverage data in normal control samples who performed analogous context updating tasks. We create support vector regression models of how target feedforward parietal-prefrontal network connectivity during context manipulation in working memory may be predicted by connectivity during simple maintenance of information, the latter process found to be less dysfunctional in schizophrenia (14-16). The machine learning models thus derived in healthy subjects were then applied to each patient to determine what his or her target “normal” quantitative feedforward connectivity should be. This novel strategy takes inspiration from related machine learning decoder approaches that utilize surrogate subjects to predict activation patterns for an individual subject with phobias, without exposure to feared stimuli (17).

## Methods

### Participants

We examined 143 healthy controls and 66 patients with schizophrenia as they performed an event-related WM task in 3T MRI scanner. All participants were enrolled as part of the Clinical Brain Disorders Branch Sibling Study (18). All participants were between 18 and 45 years of age (Table 1) and right-handed. They were interviewed by a research psychiatrist using the Structured Clinical Interview for DSM-IV, and they completed a neurological examination and a battery of neuropsychological tests. Exclusion criteria included an IQ <70, a history of prolonged substance abuse or significant medical problems, and any abnormalities found by EEG or MRI. All participants gave written consent before participation. The study was approved by the NIMH Institutional Review Board. The individuals in this sample were not part of our earlier report (1).

**Table 1.** Demographics and behavioral performance for controls and patients with schizophrenia.

|  | **CON** | **SD** | **SZ** | **SD** | **p** |
| --- | --- | --- | --- | --- | --- |
|  | (n = 143) |  | (n = 66) |  |  |
| **Sex** (number of males) | 66 |  | 44 |  | 0.01 |
| **Age** (years) | 31.027 | 9.248 | 31.909 | 10.551 | ns |
| **Parental Education** (years) | 14.23 | 3.42 | 13.82 | 3.67 | ns |
| **Education** (years) | 16.517 | 2.672 | 14.615 | 2.638 | <0.001 |
| **IQ** (WAIS) | 111.846 | 8.652 | 96.52 | 10.152 | <0.001 |
| **Accuracy** |  |  |  |  |  |
| *Working memory manipulation* | 0.933 | 0.081 | 0.776 | 0.155 | <0.001 |
| *Working memory maintenance* | 0.980 | 0.048 | 0.939 | 0.076 | <0.001 |
| **Reaction time** (s) |  |  |  |  |  |
| *Working memory manipulation* | 1.813 | 0.280 | 1.955 | 0.373 | 0.003 |
| *Working memory maintenance* | 1.039 | 0.166 | 1.181 | 0.199 | <0.001 |

CON: Controls; SZ: Patients with schizophrenia; WAIS: Wechsler Adult Intelligence Scale

### Cognitive Task Paradigm

Blood oxygen level-dependent functional MRI data were acquired from subjects after a brief training period (∼10 min). The event-related WM paradigm was described in detail in a previous study (19). In each trial, subjects first encoded 2 integer numbers presented over 1s and retained this context information in WM across a jittered interval of 3-5 seconds. In maintenance trials, subjects subsequently responded to cues as to which of the two numbers was “larger” or “smaller” within 2s. In the manipulation trials, subjects had to perform a mental subtraction of 2 or 3 on one of the two context numbers before the “larger” or “smaller” evaluation within 2s. There were 28 trials of WM manipulation and 28 trials of WM maintenance counterbalanced for trial type, and numerical size order, over about 9 minutes.

### Imaging Parameters and Analyses

T2*-weighted echo planar imaging (EPI) images with BOLD contrast were obtained with a 3T MRI scanner (GEs, Milwaukee, WI) using a standard GE head coil (64 × 64 × 24 matrix with 3.75 × 3.75 × 6.0 mm spatial resolution, repetition time (TR) =2000 ms, echo time (TE) =28 ms, flip angle=90°, field of view (FOV) =24 × 24 cm) while participants performed the cognitive task. The first four scans were discarded to allow for signal saturation.

The functional imaging data were preprocessed and analyzed using the general linear model for event-related designs in statistical parametric mapping (SPM12, Wellcome Trust Centre for Neuroimaging, London, United Kingdom). The functional images were corrected for differences in acquisition time between slices for each whole-brain volume and realigned to correct for head movement. Six movement parameters (translation: x, y, z and rotation: roll, pitch, yaw) were included in the statistical model as covariates of no interest. The functional images were normalized to a standard EPI MNI template and then spatially smoothed using an isotropic Gaussian kernel of 8 mm full-width half-maximum.

A random-effects, event-related statistical analysis was performed at two levels. In the first level, the onsets and durations of each trial for each task condition were convolved with a canonical hemodynamic response function and modeled using a general linear model on an individual subject basis. The realignment parameters were included as additional regressors of no interest. Data were high-pass filtered at 1/128 Hz. Random-effects analyses at the second (group) level were then conducted based on statistical parameter maps from each individual participant to allow population-level inference. Group level activation maps were thresholded at voxel-level whole brain p<0.05 family-wise error correction for multiple comparisons, unless otherwise stated.

### Dynamic Causal Modeling

We used Dynamic Causal Modeling, DCM (20) as implemented in SPM12 (DCM10) to examine how parietal and prefrontal brain regions interacted. We focused on the left parietal (−38 −58 38) and prefrontal (−44 30 16) regions implicated previously in context updating dysfunction in delusions (1), and examined the corresponding Human Connectome Project (21) defined parcels in the parietal (area LIPd) and prefrontal (area 46) cortex encompassing these peaks. Time series during WM context updating (manipulation) were extracted from each of the two regions-of-interest (ROI) for each individual, masked by the conjunction of the respective cortical parcel, the group-level activation mask at p<0.05 voxelwise whole brain FWE corrected for multiple comparisons, and an individual subject level activation at p<0.05. Deterministic DCM models comprising all 7 possible input and pairing combinations of these two ROIs (22) were then specified, and the strength and direction of regional interactions estimated, elucidating how regional neural activity and their interactions are influenced by cognitive inputs during WM context updating, as well as how these neuronal effects are biophysically linked to form blood-oxygen-level-dependent signals (20). Bayesian model averaging (BMA) was used to generate weighted task-related connectivity averages in each direction, for each pair of nodes, based on the posterior likelihood model fit (23, 24). Using these BMA results, we then examined task-related modulation of regional connectivity across participants, and relationships with psychopathology and polygenic risk for schizophrenia (25).

### Polygenic risk for schizophrenia

PRS for schizophrenia were calculated for each control subject as a measure of genomic risk for schizophrenia, based on the most recent GWAS (25). We obtained odds ratios of 102,217 index SNPs from a meta-analysis of PGC2 GWAS using datasets excluding the discovery sample (PGC 2014, non-lieber sample PGC2 GWAS). These 102K SNPs are LD independent (R^2^<0.1) and span across the whole genome. We then calculated a weighted sum of risk alleles for schizophrenia, by summing the imputation probability for the reference allele of the index SNP, weighted by the natural log of the odds ratio of association with schizophrenia, at each independent locus across the whole genome, as described elsewhere (25, 26). In this paper, we used the PRS calculated using the conservative PGC2 GWAS association p-value of 5e-08.

### Defining individual patient’s optimal parietal-prefrontal connectivity

We defined and tested the validity of each individual patient’s putative optimal (normal) parietal-prefrontal connectivity at WM context updating. We first created a support vector regression model, in healthy controls (N=143), of how feedforward parietal-prefrontal network connectivity during context updating may be predicted by connectivities from prefrontal and parietal regions during simple WM maintenance, a process that is less dysfunctional in schizophrenia than context updating (14-16). We included 31 parcels in the frontal and parietal cortex which were robustly engaged during WM maintenance in the patient and control samples (p<0.05 whole-brain voxel-wise FWE corrected in each separate patient or control sample). Time series during WM maintenance were then extracted from the ROI for each individual, masked by the conjunction of the respective cortical parcel, group level activation mask at p<0.05 whole brain voxelwise FWE corrected for multiple comparisons, and the individual subject level activation at p<0.05. Deterministic DCM models comprising all possible pairings across these 31 ROIs (22), giving 465 pairs of regions. From each pair of ROIs, we determined the BMA over all 7 possible models within which a pair of regions could effectively interact (as above and in (22)). This then resulted in a directed matrix of effective connectivity during WM maintenance, predicting in support vector regression modeling in healthy controls (N=143), the target parietal-prefrontal feedforward connectivity during WM context updating. We examined the in-sample predictive validity in a leave-one-out correlation analysis of the left-out predicted parietal-prefrontal connectivity versus the actual connectivity. We also created the model with half the sample and predicted the connectivity in the other half, which yielded similar results.

The support vector regression models thus derived in healthy controls were then applied to each patient to determine what that individual’s target “normal” quantitative feedforward connectivity should be at context updating during WM manipulation, given the patient’s connectivity pattern during the relatively less dysfunctional WM maintenance. We then tested the validity of each patient’s predicted normal connectivity by examining the deviation each patient’s actual connectivity was from the predicted normal, and the extent to which this deviation correlated with delusional psychopathology, previously associated with feedforward connectivity deficits. We also randomly paired each patient with a control subject, and examined the deviation between their feedforward connectivity during context updating against psychopathology, over 10,000 permutations, to test the extent to which the connectivity-derived support vector predictions may outperform that derived from simply comparing between patients and controls.

## Results

### Demographics and Behavior

Across healthy subjects (N=143) and schizophrenia patients (N=66), age and parental education were similar. Patients had a mean (±SD) duration of illness of 8.20±7.82 years with a mean age of onset 19.5±4.2years, mean total positive and negative symptom score of 54±24 and were on 298±389 mg/day chlorpromazine equivalent dose of antipsychotics. WM manipulation was more difficult (less accurate) than maintenance, and patients generally performed poorer in WM accuracy and reaction time (Table 1, p<0.001). However, WM manipulation was disproportionately more dysfunctional than WM maintenance in patients (WM process-by-diagnosis interaction F(1,217)=63.9, p<0.001, Fig 1A).

**Figure 1:**
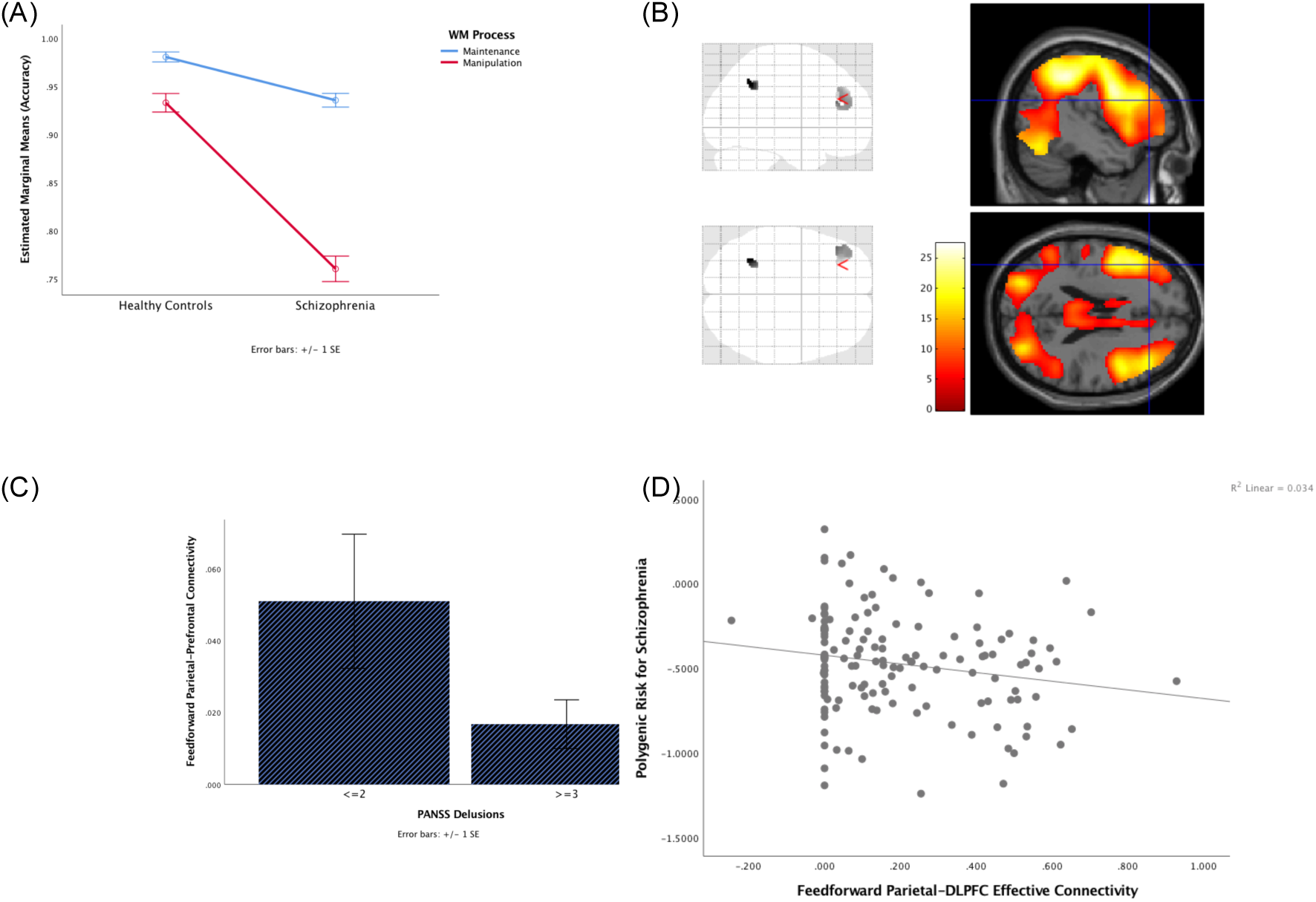
Working memory behavioral and feed-forward parietal-prefrontal connectivity. (A) Relative to controls (N=143), patients with schizophrenia (N=66) were disproportionately less accurate in performing context updating working memory manipulation, than simple maintenance of context information in WM (interaction p<0.001). (B) Patients and controls, combined (or in separate groups, not shown) robustly engaged regions-of-interest (glass-brain) in areas 46 and LIPd (p<0.05 voxelwise whole-brain FWE corrected). These ROIs encompassed regions in the DLPFC (−44 30 16) and parietal cortex (−38 −58 38) where effective connectivity during context updating was previously found to be dysfunctional in schizophrenia patients in relation to delusions (1). (C) Feedforward effective connectivity from the parietal to prefrontal cortex during context updating was relatively reduced in schizophrenia patients with delusions (N=35) vs without (N=31, p<0.05). (D) Feedforward effective connectivity from the parietal to prefrontal cortex during context updating was relatively reduced in relation to increased polygenic risk for schizophrenia in healthy controls (N=143, p<0.02).

### Parietal-prefrontal feedforward connectivity

In this report, we focus on prefrontal and parietal cortex engagement during WM. We particularly focus on the left LIPd and 46 regions-of-interest (Human Connectome Project parcellation notations (21)), which correspond to regions where we previously found parietal and prefrontal effects associated with delusions and context updating in psychosis (1). Peaks in the left LIPd and 46 ROIs were significantly engaged during context updating in WM manipulation at the group level in controls and patients (peak activation within each ROI met the activation threshold of p<0.05 voxel-wise whole-brain family-wise error FWE corrected for multiple comparisons, in each control and patient group, Supplementary Table S1). Effective connectivity across these two ROIs during context updating in WM manipulation were engaged at feedforward (parietal to prefrontal) and feedback (prefrontal to parietal) in controls (p<0.001, Supplementary Table S2) and schizophrenia patients (p<0.001, Supplementary Table S4). Connectivity in both directions was relatively reduced in patients between LIPd and 46 ROIs (p<0.001).

Within patients with schizophrenia (N=66), we examined the hypothesized role of reduced parietal-prefrontal feedforward effective connectivity at left LIPd and 46 regions during context updating at WM manipulation in delusional psychopathology suggested previously (1). Individuals with delusions on the PANSS (ratings ≥3) tended to have relatively reduced parietal-prefrontal feedforward effective connectivity (t=1.72, p<0.05; Fig 1C). Effects in the opposite feedback direction were not significant.

We then tested whether these feedforward parietal to prefrontal neural functions related to genetic mechanisms of illness rather than to confounds associated with medications or chronic ill health. If so, we would predict that there would be relationships with polygenic risk for schizophrenia in healthy individuals. Indeed, across the left parietal to dorsolateral prefrontal effective connectivity (region LIPd to 46) in healthy subjects during WM manipulation, we found reduced feedforward effective connectivity was associated with increased polygenic risk for schizophrenia (Fig 1D, r=-0.19, p<0.02). Effects in the opposite feedback direction were not significant.

### Defining individual patient’s optimal parietal-prefrontal connectivity

If indeed dysfunction in parietal-prefrontal feedforward effective connectivity during updating of new information into context may relate to delusions, the degree to which an individual patient’s connectivity may deviate from her “optimum” value should also relate to dysfunction with delusions, particularly if the connectivity optimum reflects a relevant target to normalize psychopathology. To define an individual subject’s parietal-prefrontal connectivity optimum during WM manipulation, we first examined 31 of the most activated activated parcels in the prefrontal and parietal cortices during WM maintenance in controls (peak activation in each parcel that met the activation thresholded of p<0.05 voxel-wise whole-brain family-wise error FWE corrected for multiple comparisons, Supplementary Table S3). From these brain parcels, we computed the effective connectivity from each pair of nodes in both directions, giving 930 pair-wise effective connectivity values during WM maintenance. As noted previously, WM maintenance performance accuracy was relatively less dysfunctional in patients (Cohen’s effect size d=0.673 for maintenance vs d=2.11 for manipulation, relative to controls). However, in healthy controls (N=143), within the WM maintenance prefrontal-parietal connectivity patterns in leave-one-out linear support vector regression models, feed-forward parietal-prefrontal connectivity at LIPd to 46 during WM manipulation was strongly predicted (N=143, r=0.995, p<0.00001; Fig 2A).

**Figure 2:**
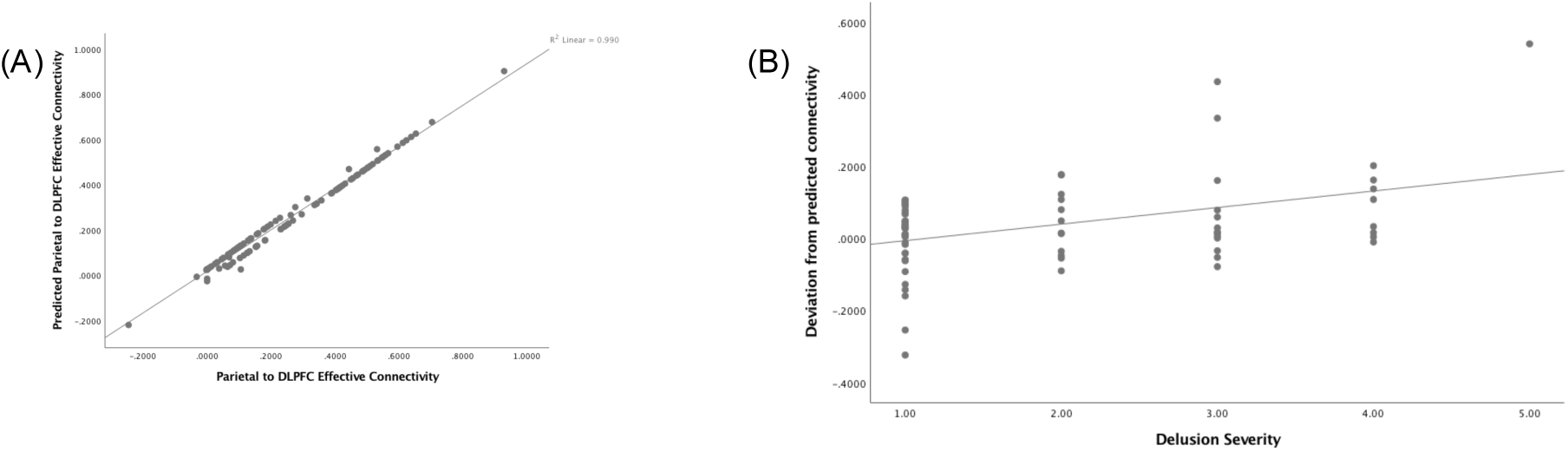
Modeled parietal-prefrontal effective connectivity. (A) Correspondence with actual effective connectivity, in healthy controls (N=143, p<0.0001), of leave-one-out predicted feedforward parietal-prefrontal effective connectivity during context updating in WM manipulation, derived from parietal and prefrontal effective connectivity patterns during WM maintenance using support vector regression models. (B) In patients with schizophrenia, deviation from predicted ‘normal’ feedforward effective connectivity correlated with delusional psychopathology (N=66, p<0.001).

We then reasoned that given WM maintenance connectivity from patients, we might predict their corresponding optimized WM manipulation connectivity, using the model derived from healthy controls. We tested the extent to which this individualized metric may be clinically significant, and found that patients’ deviations from this metric was associated with delusions (r=0.41, p<0.001; Fig 2B). On the other hand, deviation between each patient’s connectivity and randomly selected controls at parietal-prefrontal connectivity did not correlate well with psychopathology, and yielded an average r=0.01, p=0.52 over 10,000 iterations, with 2/10,000 instances of performance equal or better relative to that of the support vector machine (r>=0.41).

## Discussion

In healthy individuals, we observed that relative deficits in feedforward parietal-prefrontal connectivity during updating of new information into context during WM manipulation is associated with higher polygenic risk for schizophrenia. In schizophrenia patients, we replicate, in an independent sample, earlier observations that this parietal-prefrontal effective connectivity relates to delusional psychopathology. We then examined an estimate of optimal feedforward connectivity during WM manipulation in patients, derived from the connectivity patterns at a less dysfunctional WM maintenance task. Using a machine learning model derived from healthy subjects, larger deviations in feedforward parietal-prefrontal connectivity from predicted optima also related to delusional psychopathology.

These observations are consistent with our previous findings in a non-overlapping sample, as well as that of others on the role of feed-forward information processing deficits in perceptual and delusional psychopathology (12), and genetic risk for schizophrenia. Feed-forward parietal prefrontal connectivity during integration of new context information in WM may thus serve as a potential biomarker to uncover underlying molecular genetic mechanisms associated with information integration deficits associated with delusional psychopathology in schizophrenia. The extent to which deviation from modeled optimal connectivity may be minimized with treatment to improve psychopathology could also be examined.

Our results are consistent with a model of hierarchical posterior-cortical-to-PFC networks that integrate feed-forward information processing in order to incorporate new relevant context information (1, 6, 7, 9, 27). Dysfunction of these circuits may, in part, relate to the over-estimated likelihoods of pre-existing belief models and their over-reinforcement in delusions, and is suggested to be implicated also in the underlying genetic architecture of psychosis (1). Prefrontal dysfunction and aberrant prefrontal-parietal connectivity have also long been implicated in schizophrenia and genetic risk for psychosis (28-33). Underlying basic mechanisms of these associations could include evidence for opposing deficits in cortical dopaminergic D1 and associated glutamatergic signaling, and reciprocal exaggerated subcortical D2 signaling (34-36). These putative mechanisms have been translated through human neuroimaging studies, including ours, which also suggest interacting dopaminergic and glutamatnergic systems influencing parietal-prefrontal cortical connectivity (37), and implicate opposing D1 prefrontal-parietal and D2 prefrontal-subcortical effective connectivity in higher cognitive function dysfunctional in psychosis (38-40).

A challenge to defining individual patient targets for potential therapeutics, however, is that there is substantial inter-individual variability in cortical function, and perhaps particularly in prefrontal cortical processing (13). This would affect how one might define appropriate targets to benchmark new treatment strategies based on brain function. A promising ‘hyperalignment’ decoder approach recently utilized visual inferior temporal voxel patterns from 29 surrogate subjects to predict that for an individual with phobias without exposure to feared stimuli, the predicted data outperformed that derived at the individual-subject level (17). In terms of feedforward effective connectivity, we utilize a strategy to infer target connectivity associated with context updating based on computationally inferring a patient’s connectivity pattern from models derived from the healthy participants’ data we have collected. We infer from these data what the optimized individual patient-specific “normal” feedforward parietal-prefrontal connectivity targets would be. To the extent that patient-specific deviation from optima at this cortical feedforward connectivity may be clinically relevant, we found that these predictions do relate to psychopathology previously associated with these connections.

The specific left parietal-prefrontal feedforward connectivity target (at HCP LIPd and 46 parcels) engaged by context task stimuli similar to that proposed here has been associated with the cognitive architecture of delusions and psychosis genetic risk in independent studies (1). However, because there may be other neural connections that influence delusion formation beyond our ROIs, a larger set of brain regions and larger samples that would allow control for multiple testing should be explored. This likely would include some degree of feedback, and circular feedforward and feedback effects conceptualized in the complex neural circuit dynamics of delusions (41). Nevertheless, feedforward parietal-prefrontal connectivity during information updating appears to be at least one target associated with the cognitive and genetic underpinnings of delusional psychopathology in schizophrenia and perhaps in other psychotic conditions. The extent to which this circuit dysfunction might be causal would be important to determine, potentially with neurofeedback or other therapeutic strategies in functional MRI or EEG. Such new strategies may complement treatment of residual psychopathology, often refractory to pharmacology or cognitive-behavioral therapies.

## Acknowledgements

This work was funded by the National Institute of Mental Health Intramural Research Program (DRW, KFB), Lieber Institute for Brain Development, and National Institute of Mental Health grant R01MH101053 (HYT).

## Disclosures

None

## References

1. Kaplan C.M., et al. (2016) Estimating changing contexts in schizophrenia. Brain 139:2082–2095.

2. Woodward TS, et al. (2014) Symptom Dimensions of the Psychotic Symptom Rating Scales in Psychosis: A Multisite Study. Schizophrenia Bulletin 40(Suppl_4):S265–S274.

3. Corlett PR, Taylor JR, Wang XJ, Fletcher PC, & Krystal JH (2010) Toward a neurobiology of delusions. Prog Neurobiol 92(3):345–369.

4. Garety PA & Freeman D (1999) Cognitive approaches to delusions: a critical review of theories and evidence. The British journal of clinical psychology / the British Psychological Society 38 (Pt 2):113–154.

5. Sass L & Byrom G (2015) Phenomenological and neurocognitive perspectives on delusions: A critical overview. World Psychiatry 14(2):164–173.

6. Adams RA, Stephan KE, Brown HR, Frith CD, & Friston KJ (2013) The computational anatomy of psychosis. Frontiers in Psychiatry 4.

7. Fletcher PC & Frith CD (2009) Perceiving is believing: a Bayesian approach to explaining the positive symptoms of schizophrenia. Nat Rev Neurosci 10(1):48–58.

8. Friston K (2012) The history of the future of the Bayesian brain. Neuroimage 62(2):1230–1233.

9. Corlett PR, et al. (2007) Disrupted prediction-error signal in psychosis: evidence for an associative account of delusions. Brain 130(9):2387–2400.

10. Menon M, et al. (2011) Exploring the neural correlates of delusions of reference. Biol Psychiatry 70(12):1127–1133.

11. Teufel C, et al. (2015) Shift toward prior knowledge confers a perceptual advantage in early psychosis and psychosis-prone healthy individuals. Proceedings of the National Academy of Sciences 112(43):13401–13406.

12. Powers AR, Mathys C, & Corlett PR (2017) Pavlovian conditioning-induced hallucinations result from overweighting of perceptual priors. Science 357(6351):596–600.

13. Dubois J & Adolphs R (2016) Building a Science of Individual Differences from fMRI. Trends in Cognitive Sciences 20(6):425–443.

14. Tan HY, Choo WC, Fones CSL, & Chee MWL (2005) fMRI study of maintenance and manipulation processes within working memory in first-episode schizophrenia. American Journal of Psychiatry 162(10):1849–1858.

15. Barch DM, et al. (2001) Selective deficits in prefrontal cortex function in medication-naive patients with schizophrenia. Archives of General Psychiatry 58(3):280–288.

16. Cannon TD, et al. (2005) Dorsolateral Prefrontal Cortex Activity During Maintenance and Manipulation of Information in Working Memory in Patients With Schizophrenia. Arch Gen Psychiatry 62(10):1071–1080.

17. Taschereau-Dumouchel V, et al. (2018) Towards an unconscious neural reinforcement intervention for common fears. Proceedings of the National Academy of Sciences.

18. Egan MF, et al. (2001) Relative risk for cognitive impairments in siblings of patients with schizophrenia. Biol Psychiatry 50(2):98–107.

19. Tan HY, Callicott JH, & Weinberger DR (2007) Dysfunctional and compensatory prefrontal cortical systems, genes and the pathogenesis of schizophrenia. Cereb Cortex 17 Suppl 1:i171–181.

20. Friston KJ, Harrison L, & Penny W (2003) Dynamic causal modelling. NeuroImage 19(4):1273.

21. Glasser MF, et al. (2016) A multi-modal parcellation of human cerebral cortex. Nature 536:171.

22. Nicholson AA, et al. (2017) Dynamic causal modeling in PTSD and its dissociative subtype: Bottom–up versus top–down processing within fear and emotion regulation circuitry. Human Brain Mapping 38(11):5551–5561.

23. Penny WD, Stephan KE, Mechelli A, & Friston KJ (2004) Comparing dynamic causal models. NeuroImage 22(3):1157.

24. Stephan KE, et al. (2007) Dynamic causal models of neural system dynamics:current state and future extensions. J Biosci 32(1):129–144.

25. Psychiatric-Genetics-Consortium (2014) Biological insights from 108 schizophrenia-associated genetic loci. Nature 511(7510):421–427.

26. Consortium TIS (2009) Common polygenic variation contributes to risk of schizophrenia and bipolar disorder. Nature 460(7256):748.

27. Friston K, et al. (2016) Active inference and learning. Neuroscience & Biobehavioral Reviews 68:862–879.

28. Tan HY, Callicott JH, & Weinberger DR (2007) Dysfunctional and compensatory prefrontal cortical systems, genes and the pathogenesis of schizophrenia. Cerebral Cortex 17:i171–181.

29. Tan HY, et al. (2012) Effective connectivity of AKT1-mediated dopaminergic working memory networks and its relationship to the pharmacogenetics of cognition in schizophrenia Brain 135(5):1436–1445.

30. Tan HY, et al. (2008) Genetic variation in AKT1 is linked to dopamine-associated prefrontal cortical structure and function in humans. J Clin Invest 118(6):2200–2208.

31. Tan HY, et al. (2006) Dysfunctional prefrontal regional specialization and compensation in schizophrenia. American Journal of Psychiatry 163:1969–1977.

32. Callicott JH, et al. (2003) Abnormal fMRI Response of the Dorsolateral Prefrontal Cortex in Cognitively Intact Siblings of Patients With Schizophrenia. American Journal of Psychiatry 160(4):709–719.

33. MacDonald AW, III, et al. (2005) Specificity of Prefrontal Dysfunction and Context Processing Deficits to Schizophrenia in Never-Medicated Patients With First-Episode Psychosis. Am J Psychiatry 162(3):475–484.

34. Akil M, et al. (2003) Catechol-O-Methyltransferase Genotype and Dopamine Regulation in the Human Brain. J Neurosci 23(6):2008–2013.

35. O’Reilly RC (2006) Biologically based computational models of high-level cognition. Science 314:91–94.

36. Seamans JK & Yang CR (2004) The principal features and mechanisms of dopamine modulation in the prefrontal cortex. Prog Neurobiol 74:1–57.

37. Tan HY, et al. (2007) Epistasis between catechol-O-methyltransferase and type II metabotropic glutamate receptor 3 genes in working memory brain function. Proceedings of the National Academy of Sciences of the United States of America 104(30):12536–12541.

38. Tan HY, et al. (2012) Effective connectivity of AKT1-mediated dopaminergic working memory networks and pharmacogenetics of anti-dopaminergic treatment. Brain: a journal of neurology 135(Pt 5):1436–1445.

39. Meyer-Lindenberg A, et al. (2005) Midbrain dopamine and prefrontal function in humans: interaction and modulation by COMT genotype. Nature Neuroscience 8(5):594–596.

40. Zhang Y, et al. (2007) Polymorphisms in human dopamine D2 receptor gene affect gene expression, splicing, and neuronal activity during working memory. Proceedings of the National Academy of Sciences 104(51):20552–20557.

41. Jardri R & Denève S (2013) Circular inferences in schizophrenia. Brain 136(11):3227–3241.

