## Supplementary Tables for "Parietal–prefrontal feedforward connectivity in association with schizophrenia genetic risk and delusions"

| | SCEF | a24pr | 44–R | IFJp–R | IFSp | 46v | a1 0p | LIPd –R | FOP4 -R | AVI – R | AIP | IP2 – R | P10p | a32 pr | SFL |
|---|---|---|---|---|---|---|---|---|---|---|---|---|---|---|---|
| T-values (NC-manipulation) | 27.5 | 20.1 | 20.7 | 19.4 | 20.2 | 22.0 | 7.6 | 16.5 | 17.7 | 24.5 | 18.5 | 22.7 | 12.7 | 21.4 | 26.5 |
| T-values (NC-maintenance) | 18.9 | 15.0 | 13.9 | 12.9 | 10.5 | 11.4 | 4.6 | 15.2 | 14.9 | 17.1 | 16.7 | 17.2 | 8.3 | 14.2 | 15.6 |
| T-values (PT-manipulation) | 12.5 | 7.6 | 8.4 | 8.4 | 8.1 | 9.8 | 4.5 | 9.9 | 7.2 | 9.9 | 11.9 | 12.2 | 3.1 | 8.3 | 10.7 |
| T-values (PT-maintenance) | 14.8 | 11.5 | 7.2 | 6.2 | 5.0 | 6.0 | 4.0 | 10.6 | 7.3 | 8.0 | 11.6 | 10.4 | 2.4 | 8.3 | 10.0 |

| | 7PC | P32pr | 8BM | 44 –L | 47l | 6r | IFJa | IFJp–L | p9-46v | **46** | **LIPd–L** | FOP4 –L | AVI-L | PFt | IP2–L | FOP5 |
|---|---|---|---|---|---|---|---|---|---|---|---|---|---|---|---|---|
| T-values (NC-manipulation) | 24.7 | 21.9 | 24.7 | 23.9 | 24.0 | 28.3 | 24.9 | 23.7 | 22.3 | **14.9** | **24.61** | 19.94 | 25.8 | 26.4 | 25.9 | 21.1 |
| T-values (NC-maintenance) | 24.7 | 16.3 | 13.2 | 13.1 | 13.8 | 19.3 | 15.3 | 15.8 | 14.3 | **11.2** | **21.27** | 14.77 | 16.7 | 27.1 | 26.2 | 14.8 |
| T-values (PT-manipulation) | 13.5 | 11.2 | 9.5 | 11.7 | 7.6 | 13.1 | 11.7 | 10.4 | 9.0 | **4.9** | **12.95** | 6.91 | 8.4 | 13.5 | 13.9 | 6.9 |
| T-values (PT-maintenance) | 17.1 | 16.6 | 8.8 | 7.0 | 7.6 | 11.0 | 7.7 | 9.6 | 5.8 | **4.2** | **13.56** | 6.90 | 8.2 | 15.9 | 16.3 | 7.8 |

Supplementary Table S1: Working memory engagement in prefrontal and parietal cortices T-values of peaks in 31 most engaged regions in the parietal and prefrontal cortex in normal controls (NC, N=143), during working memory (WM) manipulation and maintenance, with corresponding engagement in schizophrenia patients (PT, N=66). All peaks survived $p<0.05$ voxelwise whole-brain family-wise (FWE) correction for multiple comparisons. Regions are annotated according to the Human Connectome Project (HCP) parcellations (1). In bold are the left 46 and LIPd regions-of.-interest corresponding to previous connectivity findings associated with dysfunctional context updating and delusions (2) that are examined in the independent samples herein.

| | SCEF | a24pr | 44 –R | IFJp –R | IFSp | 46v | a10p | LIPd –R | FOP4 -R | AVI –R | AIP | IP2 –R | P10p | a32pr | SFL | 7PC | P32pr | 8BM | 44 –L | 47l | 6r | IFJa | IFJp –L | p9-46v | **46** | **LIPd –L** | FOP4 –L | AVI –L | PFt | IP2 –L | FOP5 |
|---|---|---|---|---|---|---|---|---|---|---|---|---|---|---|---|---|---|---|---|---|---|---|---|---|---|---|---|---|---|---|---|
| SCEF | NaN | 10.7 | 16.1 | 12.8 | 13.4 | 13.4 | 6.5 | 11.9 | 14.2 | 14.9 | 13.1 | 14.6 | 7.9 | 11.1 | 15.3 | 13.7 | 12.6 | 13.4 | 14.7 | 12.8 | 15.2 | 14.3 | 14.8 | 13.1 | 10.8 | 14.3 | 14.0 | 15.5 | 11.9 | 14.8 | 12.4 |
| a24pr | 8.5 | NaN | 10.5 | 9.9 | 10.9 | 10.6 | 7.2 | 10.0 | 10.5 | 9.9 | 9.9 | 11.1 | 9.1 | 11.9 | 9.6 | 10.6 | 11.2 | 6.9 | 8.8 | 10.5 | 8.8 | 10.0 | 10.2 | 9.6 | 10.5 | 10.1 | 10.9 | 10.4 | 10.5 | 10.5 | 11.8 |
| 44 –R | 13.8 | 12.0 | NaN | 13.8 | 13.4 | 13.5 | 8.7 | 12.3 | 15.0 | 13.8 | 12.8 | 14.3 | 9.9 | 12.2 | 13.9 | 13.8 | 12.0 | 12.7 | 14.1 | 14.1 | 12.7 | 13.5 | 13.4 | 12.7 | 12.3 | 12.6 | 12.9 | 15.0 | 12.4 | 13.0 | 13.4 |
| IFJp –R | 11.0 | 10.8 | 11.2 | NaN | 11.6 | 10.8 | 7.7 | 10.3 | 11.6 | 10.6 | 11.2 | 11.0 | 10.6 | 10.5 | 10.8 | 11.1 | 10.1 | 9.9 | 10.6 | 10.4 | 9.7 | 10.7 | 10.4 | 10.5 | 11.3 | 10.6 | 11.2 | 10.0 | 9.9 | 10.8 | 11.5 |
| IFSp | 9.9 | 11.6 | 12.4 | 12.1 | NaN | 12.1 | 8.9 | 11.5 | 12.7 | 11.6 | 11.6 | 11.7 | 10.1 | 11.7 | 11.1 | 11.1 | 11.3 | 10.6 | 10.7 | 11.9 | 9.7 | 11.0 | 10.9 | 11.4 | 11.3 | 11.4 | 12.3 | 12.2 | 11.2 | 10.9 | 12.9 |
| 46v | 12.2 | 12.7 | 14.7 | 12.9 | 14.6 | NaN | 9.1 | 12.4 | 13.7 | 14.3 | 12.8 | 13.7 | 9.5 | 13.7 | 13.2 | 13.4 | 12.6 | 13.2 | 10.2 | 13.1 | 11.4 | 12.5 | 12.5 | 13.8 | 12.9 | 13.7 | 11.5 | 14.0 | 12.3 | 12.8 | 13.3 |
| a10p | 6.3 | 4.0 | 7.6 | 7.2 | 4.8 | 3.6 | NaN | 4.5 | 8.2 | 7.3 | 3.9 | 7.6 | 7.5 | 4.5 | 7.0 | 4.7 | 7.6 | 3.6 | 6.8 | 3.8 | 6.5 | 3.3 | 4.0 | 7.4 | 3.6 | 4.3 | 3.9 | 4.5 | 7.5 | 6.7 | 4.4 |
| LIPd –R | 10.0 | 9.9 | 10.4 | 10.4 | 10.2 | 9.7 | 7.7 | NaN | 11.0 | 9.8 | 11.9 | 10.7 | 9.6 | 9.8 | 9.6 | 10.3 | 9.8 | 9.7 | 9.9 | 10.6 | 8.8 | 9.6 | 9.8 | 9.7 | 10.2 | 10.2 | 10.6 | 10.5 | 10.3 | 9.8 | 10.7 |
| FOP4 -R | 10.9 | 9.6 | 11.7 | 11.3 | 11.0 | 10.9 | 7.5 | 10.9 | NaN | 11.0 | 10.4 | 11.0 | 10.4 | 10.0 | 10.6 | 10.9 | 10.5 | 9.1 | 10.9 | 11.0 | 10.5 | 10.8 | 10.2 | 10.5 | 10.5 | 10.1 | 11.9 | 11.5 | 11.2 | 10.0 | 12.3 |
| AVI –R | 12.3 | 10.7 | 13.7 | 11.9 | 12.0 | 12.5 | 8.4 | 11.4 | 14.3 | NaN | 10.6 | 11.9 | 8.4 | 11.5 | 12.4 | 12.1 | 11.8 | 11.3 | 10.0 | 12.8 | 11.4 | 12.9 | 12.7 | 12.5 | 11.6 | 12.3 | 13.9 | 14.7 | 11.8 | 12.6 | 13.5 |
| AIP | 10.8 | 10.1 | 11.1 | 11.5 | 11.0 | 10.8 | 8.2 | 13.5 | 11.6 | 10.3 | NaN | 11.7 | 8.8 | 10.7 | 9.5 | 11.5 | 10.2 | 10.1 | 10.5 | 11.1 | 10.1 | 10.1 | 10.8 | 11.0 | 10.8 | 11.5 | 11.0 | 11.2 | 11.6 | 10.8 | 12.0 |
| IP2 –R | 13.6 | 13.2 | 14.4 | 12.7 | 13.5 | 12.7 | 8.7 | 13.6 | 14.1 | 13.2 | 14.0 | NaN | 9.6 | 13.8 | 12.4 | 12.6 | 12.2 | 13.2 | 12.3 | 13.2 | 10.6 | 12.5 | 12.2 | 13.4 | 13.1 | 13.8 | 13.6 | 13.5 | 12.2 | 11.8 | 12.1 |
| P10p | 8.0 | 9.6 | 8.3 | 9.4 | 8.3 | 7.9 | 7.4 | 9.8 | 10.1 | 8.7 | 8.8 | 9.6 | NaN | 9.3 | 7.4 | 8.9 | 9.1 | 7.4 | 6.8 | 8.1 | 6.4 | 8.2 | 9.4 | 7.3 | 7.4 | 8.8 | 8.4 | 7.7 | 9.2 | 6.4 | 10.0 |
| a32pr | 8.7 | 12.6 | 10.8 | 10.9 | 11.5 | 11.1 | 8.2 | 11.1 | 11.0 | 10.6 | 10.9 | 11.4 | 9.7 | NaN | 10.1 | 11.3 | 11.2 | 9.9 | 8.8 | 10.8 | 9.1 | 9.7 | 10.3 | 10.8 | 11.4 | 10.5 | 11.3 | 11.3 | 10.7 | 10.6 | 12.4 |
| SFL | 13.6 | 11.6 | 14.4 | 12.2 | 12.1 | 13.0 | 7.8 | 11.8 | 13.4 | 13.4 | 11.6 | 13.6 | 7.9 | 12.3 | NaN | 14.0 | 12.5 | 14.4 | 13.9 | 12.0 | 14.1 | 13.6 | 14.1 | 13.6 | 11.2 | 13.9 | 13.3 | 14.2 | 12.2 | 14.0 | 12.4 |
| 7PC | 12.3 | 11.2 | 14.2 | 12.2 | 11.6 | 11.7 | 8.8 | 12.7 | 13.2 | 12.6 | 14.1 | 13.7 | 9.0 | 12.8 | 13.2 | NaN | 12.5 | 12.2 | 13.1 | 11.9 | 12.6 | 13.3 | 12.6 | 11.3 | 14.3 | 14.5 | 13.7 | 13.2 | 13.0 | 12.7 | 8.0 |
| P32pr | 10.2 | 12.4 | 11.6 | 11.1 | 11.5 | 11.0 | 7.5 | 11.0 | 11.8 | 11.1 | 10.8 | 11.9 | 9.7 | 11.6 | 10.8 | 13.7 | NaN | 10.5 | 11.2 | 10.7 | 10.8 | 11.0 | 10.6 | 10.2 | 11.2 | 10.2 | 11.6 | 10.6 | 11.0 | 11.0 | 11.1 |
| 8BM | 12.9 | 10.9 | 13.5 | 11.9 | 12.0 | 13.4 | 7.8 | 11.8 | 12.8 | 12.5 | 12.2 | 13.7 | 7.5 | 12.6 | 14.4 | 13.0 | 12.7 | NaN | 12.7 | 12.9 | 11.6 | 12.2 | 13.5 | 13.6 | 10.8 | 13.3 | 12.7 | 13.7 | 12.3 | 13.6 | 12.2 |
| 44 –L | 14.1 | 11.1 | 15.4 | 13.4 | 13.1 | 13.5 | 8.0 | 12.2 | 14.5 | 15.7 | 13.1 | 13.5 | 8.6 | 12.0 | 9.4 | 12.4 | 11.3 | 13.3 | NaN | 11.6 | 13.3 | 14.3 | 13.5 | 13.3 | 11.6 | 13.5 | 14.1 | 11.8 | 13.1 | 13.9 | 14.0 |
| 47l | 10.6 | 11.6 | 12.5 | 11.4 | 12.1 | 11.2 | 8.2 | 11.4 | 13.1 | 11.3 | 11.2 | 11.6 | 8.8 | 9.7 | 10.1 | 13.6 | 12.3 | 11.3 | 11.5 | NaN | 11.3 | 11.2 | 12.8 | 11.7 | 9.3 | 11.8 | 13.8 | 13.7 | 12.5 | 10.9 | 12.3 |
| 6r | 15.5 | 10.5 | 15.9 | 12.8 | 12.4 | 12.9 | 7.1 | 11.6 | 13.5 | 13.9 | 11.8 | 12.5 | 7.3 | 11.7 | 14.8 | 13.2 | 11.9 | 13.1 | 14.2 | 13.2 | NaN | 14.2 | 15.5 | 11.5 | 14.0 | 15.3 | 14.7 | 13.2 | 14.8 | 11.3 | 11.3 |
| IFJa | 13.0 | 11.3 | 14.5 | 13.6 | 13.1 | 12.6 | 7.3 | 11.6 | 14.4 | 14.0 | 12.1 | 13.3 | 8.9 | 11.5 | 13.6 | 13.3 | 11.9 | 12.5 | 13.6 | 13.5 | 12.1 | NaN | 12.9 | 13.1 | 12.0 | 14.0 | 13.2 | 14.0 | 12.8 | 13.6 | 13.6 |
| IFJp –L | 13.4 | 11.9 | 14.3 | 12.9 | 12.0 | 12.6 | 8.6 | 12.2 | 13.0 | 13.2 | 12.0 | 12.6 | 9.4 | 12.1 | 13.3 | 13.0 | 11.5 | 12.8 | 12.3 | 13.5 | 12.8 | 12.4 | NaN | 13.3 | 12.4 | 14.7 | 13.8 | 14.3 | 12.2 | 12.2 | 12.6 |
| p9-46v | 12.0 | 11.3 | 13.8 | 13.2 | 13.3 | 13.6 | 8.0 | 11.6 | 13.1 | 13.1 | 12.6 | 13.4 | 9.6 | 12.1 | 13.1 | 11.1 | 10.6 | 12.9 | 9.1 | 13.1 | 8.7 | 12.3 | 12.4 | NaN | 12.1 | 12.9 | 12.9 | 13.6 | 13.5 | 13.0 | 12.1 |
| **46** | 9.4 | 11.1 | 11.5 | 11.6 | 11.8 | 11.1 | 8.3 | 10.9 | 12.1 | 11.0 | 11.0 | 11.0 | 9.9 | 11.1 | 10.5 | 13.4 | 11.4 | 10.4 | 9.8 | 11.0 | 11.0 | 10.1 | 9.7 | 9.8 | NaN | **10.7** | 10.1 | 11.7 | 10.6 | 10.6 | 12.2 |
| **LIPd –L** | 12.2 | 11.1 | 11.9 | 12.5 | 11.9 | 12.1 | 8.6 | 12.1 | 12.9 | 13.2 | 12.4 | 13.2 | 10.1 | 11.8 | 12.3 | 15.0 | 12.2 | 11.8 | 11.5 | 12.5 | 13.1 | 12.0 | 12.1 | 12.3 | **12.2** | NaN | 12.7 | 12.8 | 12.7 | 12.0 | 12.5 |

| | | | | | | | | | | | | | | | | | | | | | | | | | | | | | | | |
|---|---|---|---|---|---|---|---|---|---|---|---|---|---|---|---|---|---|---|---|---|---|---|---|---|---|---|---|---|---|---|---|
| FOP4 –L | 10.9 | 10.7 | 11.1 | 10.7 | 11.8 | 10.8 | 7.9 | 11.3 | 13.2 | 11.8 | 10.1 | 11.3 | 8.3 | 11.4 | 11.0 | 12.2 | 11.8 | 10.1 | 10.6 | 12.7 | 11.3 | 10.9 | 11.6 | 11.3 | 9.9 | 11.3 | NaN | 12.3 | 10.7 | 10.5 | 12.9 |
| AVI –L | 11.4 | 11.4 | 13.1 | 11.1 | 12.7 | 12.0 | 8.5 | 11.6 | 13.5 | 13.9 | 11.3 | 12.4 | 9.0 | 12.5 | 11.7 | 13.6 | 11.8 | 11.8 | 12.2 | 13.9 | 12.1 | 12.0 | 12.4 | 12.2 | 12.2 | 12.1 | 14.5 | NaN | 11.8 | 11.8 | 13.6 |
| PFt | 11.5 | 11.7 | 12.0 | 11.2 | 11.6 | 10.9 | 8.3 | 12.2 | 13.4 | 11.5 | 13.4 | 12.1 | 10.0 | 12.8 | 11.3 | 15.4 | 12.2 | 11.9 | 11.6 | 13.5 | 13.7 | 12.0 | 11.4 | 13.0 | 11.3 | 12.7 | 12.1 | 12.2 | NaN | 10.8 | 13.9 |
| IP2 –L | 14.6 | 12.1 | 15.1 | 13.5 | 12.6 | 13.8 | 7.8 | 12.9 | 13.8 | 15.2 | 13.6 | 14.9 | 8.8 | 13.2 | 15.0 | 10.6 | 12.2 | 14.7 | 12.2 | 14.1 | 8.5 | 13.6 | 13.6 | 14.7 | 12.8 | 15.2 | 14.0 | 15.3 | 13.6 | NaN | 13.6 |
| FOP5 | 9.3 | 10.6 | 10.0 | 10.4 | 10.5 | 10.1 | 8.1 | 9.9 | 11.2 | 10.2 | 9.7 | 10.1 | 9.4 | 11.0 | 9.6 | 11.1 | 8.9 | 8.5 | 9.4 | 11.4 | 8.5 | 10.1 | 10.0 | 9.5 | 10.1 | 10.2 | 11.3 | 10.9 | 10.7 | 9.9 | NaN |

Supplementary Table S2: Effective connectivity (DCM) for controls (N=143) during WM manipulation: T-values of effective connectivity across the 31 cortical parcels most engaged during WM manipulation. Rows represent connectivity “to” that region, while columns represent connectivity “from” that region. Regions are annotated according to the Human Connectome Project (HCP) parcellations(1). In bold are the left 46 and LIPd regions-of.-interest corresponding to previous connectivity findings associated with dysfunctional context updating and delusions(2) that are examined in the independent samples herein.

| | SCEF | a24pr | 44 – R | IFJp –R | IFSp | 46v | a10p | LIPd –R | FOP4 -R | AVI –R | AIP | IP2 – R | P10p | a32pr | SFL | 7PC | P32pr | 8BM | 44 – L | 47l | 6r | IFJa | IFJp –L | p9-46v | 46 | LIPd –L | FOP4 –L | AVI –L | PFt | IP2 – L | FOP5 |
|---|---|---|---|---|---|---|---|---|---|---|---|---|---|---|---|---|---|---|---|---|---|---|---|---|---|---|---|---|---|---|---|
| SCEF | NaN | 9.2 | 3.8 | 4.5 | 6.6 | 8.3 | 4.1 | 8.1 | 8.6 | 7.7 | 9.3 | 9.7 | 6.9 | 8.1 | 8.5 | 9.3 | 9.6 | 5.7 | 9.5 | 7.7 | 6.3 | 7.8 | 7.0 | 9.0 | 9.1 | 8.2 | 10.0 | 8.4 | 8.9 | 10.0 | 9.8 |
| a24pr | 8.0 | NaN | 3.3 | 4.7 | 6.4 | 5.6 | 4.4 | 7.5 | 7.2 | 8.4 | 7.8 | 8.3 | 5.0 | 7.2 | 7.3 | 8.2 | 6.9 | 6.4 | 6.9 | 7.3 | 7.2 | 7.3 | 6.7 | 5.6 | 6.9 | 8.0 | 8.6 | 7.2 | 6.6 | 8.4 | 3.6 |
| 44 –R | 6.5 | 6.7 | NaN | 3.9 | 4.5 | 5.5 | 4.7 | 8.3 | 4.7 | 7.4 | 8.2 | 8.5 | 7.4 | 7.8 | 6.2 | 7.4 | 4.6 | 6.9 | 8.8 | 7.2 | 6.3 | 6.3 | 4.3 | 6.6 | 8.0 | 7.7 | 9.1 | 6.5 | 7.8 | 7.8 | 7.9 |
| IFJp – R | 1.9 | 3.8 | 3.5 | NaN | 3.6 | 3.9 | 2.7 | 4.5 | 5.4 | 2.7 | 4.1 | 5.4 | 1.6 | 1.6 | 1.9 | 4.3 | 4.2 | 3.8 | 4.5 | 2.9 | 2.0 | 1.1 | 5.9 | 5.8 | 4.2 | 3.5 | 3.1 | 4.4 | 3.1 | 4.1 | 5.8 |
| IFSp | 6.5 | 5.9 | 7.0 | 1.6 | NaN | 6.5 | 4.8 | 5.8 | 6.2 | 5.1 | 4.5 | 7.2 | 6.7 | 5.0 | 6.1 | 6.9 | 6.4 | 6.3 | 3.4 | 4.2 | 5.4 | 5.9 | 6.3 | 4.1 | 6.8 | 6.3 | 5.9 | 4.0 | 6.5 | 6.8 | 7.0 |
| 46v | 8.0 | 6.6 | 8.6 | 5.9 | 8.3 | NaN | 5.8 | 8.2 | 8.0 | 5.3 | 9.5 | 8.9 | 7.8 | 5.5 | 7.9 | 8.5 | 7.3 | 7.8 | 7.0 | 5.0 | 5.8 | 8.4 | 9.0 | 8.5 | 9.2 | 8.7 | 6.9 | 7.6 | 8.6 | 8.3 | 7.3 |
| a10p | 3.6 | 4.5 | 5.1 | 4.1 | 5.6 | 5.6 | NaN | 5.8 | 4.8 | 6.1 | 5.9 | 5.8 | 6.3 | 4.8 | 5.1 | 5.9 | 5.8 | 4.9 | 5.0 | 5.5 | 4.5 | 5.1 | 5.1 | 5.4 | 6.3 | 5.7 | 5.3 | 4.9 | 6.1 | 5.8 | 5.7 |
| LIPd –R | 4.9 | 6.6 | 3.7 | 4.2 | 6.8 | 7.4 | 5.2 | NaN | 7.6 | 6.8 | 5.7 | 7.9 | 6.3 | 6.0 | 5.2 | 7.0 | 7.3 | 5.8 | 6.9 | 7.9 | 5.5 | 5.3 | 5.0 | 7.5 | 6.8 | 6.2 | 7.3 | 7.2 | 4.0 | 7.2 | 7.8 |
| FOP4 -R | 7.0 | 8.2 | 7.7 | 1.9 | 6.2 | 6.5 | 3.9 | 8.2 | NaN | 2.9 | 2.8 | 2.8 | 6.6 | 7.6 | 5.9 | 2.6 | 5.9 | 4.6 | 7.6 | 6.9 | 4.9 | 4.9 | 6.3 | 6.4 | 3.7 | 8.0 | 3.1 | 8.4 | 7.8 | 2.8 | 3.5 |
| AVI – R | 8.6 | 7.9 | 8.3 | 2.2 | 6.1 | 4.7 | 5.9 | 8.3 | 9.8 | NaN | 7.1 | 7.9 | 6.3 | 7.1 | 4.9 | 7.7 | 7.8 | 6.1 | 6.1 | 7.5 | 5.7 | 2.8 | 5.8 | 7.9 | 7.7 | 4.7 | 8.9 | 5.8 | 7.8 | 5.4 | 9.1 |
| AIP | 6.1 | 7.4 | 5.2 | 5.3 | 7.6 | 8.8 | 5.3 | 9.8 | 7.2 | 7.1 | NaN | 9.2 | 7.1 | 3.5 | 6.0 | 8.4 | 8.2 | 3.1 | 6.6 | 7.9 | 3.3 | 2.8 | 7.1 | 8.7 | 7.4 | 7.9 | 8.2 | 7.5 | 7.8 | 7.7 | 8.2 |
| IP2 – R | 6.5 | 7.7 | 8.5 | 3.6 | 3.9 | 8.5 | 4.2 | 9.8 | 6.8 | 8.0 | 4.5 | NaN | 2.8 | 3.2 | 5.7 | 8.3 | 8.2 | 3.1 | 5.8 | 8.2 | 4.3 | 2.5 | 6.6 | 4.9 | 5.0 | 4.1 | 8.2 | 5.9 | 8.9 | 7.8 | 8.8 |
| P10p | 5.9 | 4.1 | 6.6 | 5.2 | 7.1 | 6.8 | 6.2 | 7.1 | 6.3 | 6.3 | 7.3 | 6.8 | NaN | 5.5 | 3.2 | 6.3 | 6.4 | 4.7 | 6.2 | 5.8 | 4.6 | 7.4 | 7.3 | 7.2 | 7.8 | 6.1 | 6.9 | 5.7 | 6.5 | 6.3 | 2.3 |
| a32pr | 6.5 | 6.2 | 7.8 | 5.7 | 3.1 | 5.6 | 4.8 | 6.7 | 5.5 | 5.9 | 7.1 | 7.6 | 5.6 | NaN | 6.7 | 7.6 | 6.4 | 7.4 | 5.7 | 5.6 | 5.6 | 6.8 | 6.2 | 6.3 | 7.4 | 5.7 | 7.6 | 6.5 | 7.2 | 7.1 | 3.1 |
| SFL | 8.1 | 6.1 | 7.4 | 2.5 | 6.3 | 6.5 | 5.3 | 6.8 | 3.7 | 6.1 | 8.1 | 5.9 | 6.2 | 6.4 | NaN | 8.0 | 7.3 | 4.8 | 7.6 | 7.0 | 4.6 | 6.6 | 6.2 | 7.2 | 8.0 | 4.2 | 7.3 | 6.1 | 8.5 | 7.5 | 7.7 |
| 7PC | 9.2 | 9.9 | 7.1 | 6.4 | 9.4 | 10.7 | 6.8 | 9.0 | 8.3 | 10.0 | 8.4 | 11.4 | 7.9 | 8.4 | 9.3 | NaN | 8.6 | 8.5 | 8.6 | 9.4 | 9.7 | 8.7 | 8.2 | 9.9 | 7.9 | 11.3 | 8.7 | 8.3 | 10.1 | 12.2 | 7.3 |
| P32pr | 7.2 | 6.7 | 3.1 | 3.6 | 4.4 | 6.4 | 5.7 | 7.9 | 6.4 | 3.8 | 9.0 | 8.8 | 6.1 | 7.1 | 6.0 | 7.4 | NaN | 5.9 | 8.2 | 7.7 | 6.3 | 4.4 | 5.0 | 7.1 | 7.6 | 7.1 | 8.1 | 4.6 | 7.0 | 7.5 | 6.3 |
| 8BM | 7.8 | 6.5 | 8.4 | 1.6 | 3.4 | 7.5 | 3.7 | 6.8 | 3.9 | 4.4 | 7.1 | 7.9 | 5.0 | 5.0 | 7.6 | 7.3 | 6.1 | NaN | 6.9 | 6.7 | 7.8 | 5.3 | 2.8 | 4.0 | 6.9 | 5.7 | 7.1 | 6.4 | 8.2 | 6.4 | 7.5 |
| 44 –L | 8.3 | 7.4 | 9.0 | 5.1 | 6.2 | 8.0 | 4.9 | 3.8 | 6.8 | 6.7 | 3.5 | 3.9 | 6.5 | 3.4 | 4.8 | 6.8 | 6.8 | 3.2 | NaN | 8.9 | 6.7 | 6.5 | 7.5 | 8.3 | 7.2 | 3.9 | 8.6 | 6.8 | 7.5 | 3.6 | 8.0 |
| 47l | 7.5 | 7.5 | 8.3 | 4.1 | 5.3 | 5.4 | 5.5 | 6.7 | 7.5 | 8.4 | 7.0 | 8.1 | 6.0 | 7.1 | 7.6 | 7.3 | 7.3 | 6.5 | 8.6 | NaN | 6.6 | 7.5 | 7.0 | 7.4 | 7.0 | 7.1 | 9.3 | 6.7 | 8.0 | 5.3 | 8.4 |
| 6r | 6.0 | 5.8 | 6.5 | 4.5 | 3.8 | 7.2 | 3.5 | 7.0 | 4.7 | 6.1 | 5.8 | 6.7 | 5.3 | 3.1 | 5.5 | 7.2 | 5.5 | 3.2 | 8.3 | 8.5 | NaN | 8.3 | 6.7 | 4.9 | 8.5 | 5.3 | 7.7 | 4.4 | 6.4 | 6.0 | 7.8 |
| IFJa | 5.7 | 6.3 | 5.2 | 1.6 | 7.2 | 7.2 | 3.9 | 6.0 | 4.9 | 4.8 | 6.2 | 7.6 | 3.0 | 3.1 | 0.8 | 6.6 | 4.0 | 1.3 | 7.8 | 6.9 | 3.0 | NaN | 7.5 | 3.7 | 7.8 | 5.8 | 7.1 | 5.8 | 6.6 | 5.6 | 7.3 |
| IFJp – L | 3.9 | 5.7 | 4.3 | 3.0 | 6.1 | 6.5 | 3.5 | 6.7 | 6.3 | 4.5 | 5.7 | 7.3 | 2.7 | 6.1 | 2.3 | 4.9 | 5.1 | 2.9 | 5.1 | 6.7 | 2.0 | 3.4 | NaN | 7.3 | 6.1 | 5.1 | 4.3 | 6.2 | 4.5 | 6.1 | 7.2 |

| | | | | | | | | | | | | | | | | | | | | | | | | | | | | | | | |
|---|---|---|---|---|---|---|---|---|---|---|---|---|---|---|---|---|---|---|---|---|---|---|---|---|---|---|---|---|---|---|---|
| p9-46v | 5.9 | 7.9 | 8.3 | 7.6 | 7.9 | 8.2 | 5.3 | 8.2 | 6.9 | 8.5 | 7.9 | 8.7 | 7.6 | 7.7 | 6.7 | 7.7 | 6.7 | 6.2 | 7.5 | 7.6 | 6.9 | 7.1 | 8.7 | NaN | 8.9 | 7.2 | 8.4 | 7.8 | 10.0 | 9.1 | 8.3 |
| 46 | 7.3 | 7.0 | 7.4 | 2.4 | 7.5 | 8.3 | 6.1 | 8.1 | 7.8 | 7.9 | 3.6 | 3.2 | 7.7 | 7.9 | 7.6 | 3.1 | 6.5 | 7.6 | 8.3 | 8.2 | 8.2 | 7.5 | 7.5 | 9.0 | NaN | 7.9 | 9.0 | 3.4 | 9.4 | 8.5 | 8.3 |
| LIPd –L | 7.9 | 7.9 | 9.1 | 6.5 | 7.9 | 9.6 | 6.1 | 8.7 | 8.4 | 5.6 | 8.6 | 9.1 | 7.0 | 3.8 | 6.5 | 10.0 | 8.2 | 5.3 | 8.3 | 8.1 | 5.0 | 3.4 | 6.6 | 6.5 | 6.7 | NaN | 8.6 | 6.0 | 7.3 | 8.5 | 7.8 |
| FOP4 –L | 9.3 | 3.9 | 3.6 | 3.2 | 7.9 | 2.9 | 1.8 | 4.3 | 8.6 | 5.4 | 4.1 | 3.9 | 6.2 | 3.9 | 3.3 | 3.1 | 8.0 | 3.7 | 3.4 | 8.8 | 3.3 | 7.1 | 3.9 | 4.2 | 4.4 | 3.6 | NaN | 4.5 | 4.3 | 3.6 | 4.9 |
| AVI – L | 6.1 | 6.4 | 8.6 | 4.7 | 5.9 | 6.0 | 4.2 | 7.4 | 8.4 | 8.4 | 6.5 | 7.1 | 4.6 | 6.9 | 6.0 | 5.6 | 6.7 | 5.3 | 7.6 | 6.0 | 4.3 | 5.8 | 7.6 | 8.2 | 6.7 | 6.2 | 9.1 | NaN | 5.7 | 6.5 | 9.0 |
| PFt | 8.2 | 8.2 | 7.4 | 5.7 | 8.9 | 4.2 | 6.8 | 8.8 | 8.5 | 5.0 | 5.3 | 9.2 | 7.3 | 4.9 | 8.6 | 6.5 | 8.6 | 7.0 | 7.1 | 9.6 | 7.5 | 7.2 | 6.3 | 10.0 | 10.4 | 10.9 | 10.8 | 4.8 | NaN | 12.0 | 8.3 |
| IP2 – L | 9.4 | 9.1 | 7.2 | 6.5 | 8.3 | 9.4 | 6.8 | 10.6 | 8.4 | 7.1 | 8.2 | 10.3 | 7.7 | 5.4 | 8.8 | 11.8 | 8.6 | 6.3 | 9.2 | 7.6 | 8.4 | 7.4 | 8.4 | 8.7 | 8.0 | 9.6 | 8.5 | 8.4 | 9.6 | NaN | 9.9 |
| FOP5 | 7.4 | 6.9 | 8.0 | 5.5 | 7.0 | 6.5 | 5.1 | 7.6 | 7.3 | 7.8 | 2.4 | 7.8 | 5.4 | 3.4 | 7.0 | 2.0 | 7.5 | 2.3 | 7.0 | 8.1 | 7.2 | 6.9 | 7.6 | 7.9 | 7.0 | 7.0 | 9.8 | 8.5 | 5.6 | 8.1 | NaN |

Supplementary Table S3: Effective connectivity (DCM) for controls (N=143) during WM maintenance: T-values of effective connectivity across the 31 cortical parcels most engaged during WM maintenance. Rows represent connectivity "to" that region, while columns represent connectivity "from" that region. Regions are annotated according to the Human Connectome Project (HCP) parcellations (1).

| | SCEF | a24pr | 44 –R | IFJp –R | IFSp | 46v | a10p | LIPd –R | FOP4 -R | AVI –R | AIP | IP2 –R | P10p | a32pr | SFL | 7PC | P32pr | 8BM | 44 –L | 47l | 6r | IFJa | IFJp –L | p9-46v | **46** | **LIPd –L** | FOP4 –L | AVI –L | PFt | IP2 –L | FOP5 |
|---|---|---|---|---|---|---|---|---|---|---|---|---|---|---|---|---|---|---|---|---|---|---|---|---|---|---|---|---|---|---|---|
| SCEF | NaN | 3.8 | 3.5 | 4.2 | 3.9 | 4.2 | 3.6 | 4.0 | 4.5 | 5.2 | 4.7 | 4.4 | 3.6 | 4.6 | 2.9 | 4.5 | 4.1 | 4.6 | 4.4 | 3.2 | 4.7 | 4.1 | 3.6 | 4.7 | 5.0 | 4.5 | 4.5 | 4.7 | 4.6 | 3.8 | 4.8 |
| a24pr | 3.8 | NaN | 2.2 | 2.9 | 2.9 | 2.2 | 2.8 | 3.6 | 4.3 | 4.0 | 4.0 | 2.7 | 3.6 | 4.5 | 2.3 | 3.0 | 4.7 | 3.5 | 3.3 | 3.1 | 2.0 | 2.7 | 3.7 | 3.6 | 4.7 | 3.5 | 4.5 | 4.6 | 4.5 | 2.8 | 4.3 |
| 44 –R | 3.6 | 2.4 | NaN | 3.7 | 4.2 | 4.4 | 3.4 | 3.5 | 2.4 | 4.6 | 3.3 | 5.1 | 4.1 | 3.2 | 4.4 | 4.4 | 1.6 | 3.3 | 4.2 | 3.1 | 5.1 | 4.1 | 4.1 | 3.6 | 3.2 | 4.6 | 2.5 | 3.0 | 4.0 | 4.6 | 3.5 |
| IFJp – R | 4.4 | 3.0 | 3.4 | NaN | 3.3 | 3.3 | 3.0 | 3.3 | 3.8 | 3.7 | 4.4 | 4.8 | 3.5 | 3.4 | 3.2 | 4.0 | 2.9 | 3.0 | 3.2 | 3.8 | 3.9 | 3.8 | 3.5 | 3.3 | 3.5 | 3.9 | 3.9 | 4.5 | 3.7 | 4.4 | 3.9 |
| IFSp | 3.7 | 2.7 | 4.4 | 3.5 | NaN | 4.7 | 3.2 | 3.5 | 2.9 | 3.7 | 3.8 | 4.5 | 4.0 | 2.8 | 3.3 | 3.9 | 2.7 | 2.9 | 4.4 | 3.9 | 4.2 | 4.1 | 3.9 | 3.6 | 2.4 | 4.0 | 2.7 | 3.5 | 2.8 | 4.2 | 2.9 |
| 46v | 4.0 | 2.8 | 4.5 | 3.7 | 5.4 | NaN | 4.3 | 3.5 | 2.6 | 4.5 | 3.8 | 4.9 | 4.0 | 2.4 | 3.5 | 5.4 | 2.4 | 4.1 | 4.9 | 2.9 | 4.8 | 4.4 | 4.4 | 4.6 | 3.7 | 3.9 | 2.7 | 3.7 | 3.2 | 4.5 | 2.4 |
| a10p | 3.4 | 2.6 | 3.6 | 3.4 | 4.0 | 4.4 | NaN | 3.0 | 2.5 | 4.0 | 2.9 | 4.1 | 3.8 | 2.5 | 3.0 | 3.1 | 2.4 | 3.9 | 4.0 | 2.8 | 3.7 | 3.1 | 3.6 | 1.9 | 3.0 | 3.6 | 2.6 | 3.1 | 2.2 | 3.6 | 2.5 |
| LIPd –R | 4.4 | 3.5 | 3.6 | 3.7 | 3.8 | 3.8 | 2.8 | NaN | 3.7 | 4.2 | 3.8 | 4.4 | 3.4 | 3.8 | 3.4 | 4.0 | 3.3 | 3.6 | 3.8 | 3.4 | 3.6 | 4.4 | 4.2 | 3.5 | 4.1 | 4.4 | 4.5 | 4.2 | 3.9 | 4.3 | 3.6 |
| FOP4 -R | 4.6 | 3.8 | 1.7 | 4.4 | 2.8 | 3.1 | 1.5 | 4.0 | NaN | 4.5 | 4.4 | 3.5 | 3.0 | 4.0 | 2.5 | 3.2 | 4.3 | 1.6 | 3.4 | 3.8 | 2.7 | 3.0 | 3.9 | 3.1 | 4.0 | 4.1 | 4.5 | 4.3 | 4.1 | 2.7 | 4.8 |
| AVI – R | 5.0 | 3.9 | 4.7 | 4.2 | 4.1 | 4.8 | 3.6 | 4.0 | 3.7 | NaN | 4.6 | 5.6 | 4.0 | 3.9 | 3.5 | 4.7 | 3.8 | 4.5 | 4.6 | 4.2 | 4.8 | 4.3 | 4.1 | 4.2 | 4.8 | 3.9 | 3.8 | 4.4 | 4.3 | 3.5 | 4.6 |
| AIP | 5.0 | 4.1 | 3.5 | 4.4 | 3.9 | 4.2 | 2.8 | 3.9 | 4.2 | 4.9 | NaN | 4.4 | 3.5 | 4.3 | 3.4 | 5.2 | 3.6 | 3.1 | 4.1 | 3.9 | 3.7 | 4.4 | 4.4 | 3.7 | 4.3 | 4.8 | 4.6 | 4.3 | 4.3 | 4.8 | 4.3 |
| IP2 – R | 4.0 | 2.7 | 4.8 | 4.0 | 4.2 | 4.8 | 4.0 | 3.9 | 3.3 | 4.5 | 4.4 | NaN | 3.8 | 2.7 | 3.4 | 5.0 | 2.7 | 4.6 | 4.9 | 3.9 | 4.7 | 4.7 | 4.2 | 4.5 | 3.6 | 4.6 | 3.6 | 3.8 | 4.0 | 5.0 | 3.9 |
| P10p | 3.8 | 3.1 | 3.6 | 4.4 | 4.7 | 4.3 | 3.4 | 3.7 | 3.0 | 4.2 | 4.1 | 3.9 | NaN | 3.1 | 3.7 | 2.6 | 2.8 | 3.5 | 3.7 | 3.4 | 4.2 | 4.0 | 4.4 | 2.6 | 3.0 | 4.1 | 3.4 | 4.0 | 3.9 | 3.6 | 3.5 |
| a32pr | 4.5 | 4.9 | 3.5 | 3.7 | 3.0 | 2.4 | 2.6 | 4.1 | 4.2 | 3.8 | 4.4 | 3.1 | 3.6 | NaN | 3.1 | 3.9 | 4.8 | 3.7 | 4.3 | 4.2 | 2.8 | 4.2 | 3.6 | 4.1 | 4.0 | 4.0 | 4.7 | 4.6 | 4.3 | 3.1 | 4.3 |
| SFL | 3.0 | 3.0 | 4.4 | 3.3 | 3.8 | 3.7 | 2.4 | 3.4 | 3.0 | 3.5 | 3.7 | 4.1 | 3.4 | 3.2 | NaN | 4.0 | 2.9 | 3.6 | 4.3 | 3.5 | 4.6 | 3.5 | 3.9 | 2.9 | 2.2 | 3.9 | 3.4 | 3.0 | 3.0 | 3.3 | 2.6 |
| 7PC | 4.8 | 3.3 | 4.2 | 3.9 | 4.7 | 5.2 | 2.9 | 4.6 | 3.4 | 4.7 | 4.8 | 5.3 | 3.1 | 3.3 | 4.0 | NaN | 2.5 | 3.5 | 4.7 | 4.2 | 3.9 | 4.0 | 4.3 | 3.3 | 2.7 | 5.8 | 3.6 | 4.2 | 3.0 | 5.9 | 3.2 |
| P32pr | 4.4 | 5.1 | 2.3 | 2.8 | 1.9 | 2.0 | 2.2 | 3.9 | 4.8 | 4.1 | 3.8 | 2.8 | 3.0 | 4.5 | 1.9 | 2.6 | NaN | 3.4 | 3.5 | 2.8 | 2.1 | 3.0 | 3.5 | 3.8 | 4.6 | 3.4 | 4.6 | 4.6 | 4.1 | 2.5 | 4.5 |
| 8BM | 4.0 | 3.0 | 4.0 | 3.2 | 3.4 | 3.7 | 3.6 | 3.8 | 2.2 | 4.0 | 3.6 | 4.2 | 3.8 | 3.6 | 3.4 | 3.6 | 3.1 | NaN | 4.4 | 2.9 | 4.3 | 4.5 | 4.3 | 4.1 | 3.7 | 3.9 | 2.9 | 3.2 | 3.3 | 4.1 | 2.5 |
| 44 –L | 4.1 | 2.8 | 4.5 | 3.3 | 4.3 | 4.7 | 1.6 | 3.7 | 3.6 | 4.5 | 4.2 | 5.3 | 3.2 | 4.0 | 4.1 | 5.3 | 2.4 | 3.5 | NaN | 4.3 | 5.0 | 5.1 | 5.0 | 3.3 | 2.5 | 5.1 | 3.6 | 2.8 | 2.9 | 5.0 | 3.9 |
| 47l | 3.3 | 3.3 | 3.1 | 4.0 | 4.0 | 3.1 | 3.3 | 3.6 | 4.0 | 4.3 | 3.6 | 3.7 | 3.0 | 4.2 | 2.9 | 4.0 | 3.4 | 3.1 | 3.7 | NaN | 4.5 | 3.8 | 4.5 | 3.6 | 3.7 | 4.5 | 4.1 | 4.4 | 4.3 | 3.2 | 4.3 |
| 6r | 4.7 | 2.2 | 5.3 | 3.3 | 4.7 | 5.3 | 3.9 | 4.1 | 3.1 | 4.6 | 4.2 | 5.3 | 3.6 | 3.1 | 3.8 | 4.1 | 2.6 | 4.4 | 5.7 | 4.2 | NaN | 4.7 | 4.7 | 3.5 | 2.2 | 4.7 | 2.9 | 3.7 | 3.1 | 5.0 | 3.1 |
| IFJa | 4.0 | 3.3 | 5.0 | 4.4 | 4.4 | 5.0 | 3.3 | 3.4 | 3.3 | 4.4 | 4.0 | 5.4 | 3.8 | 3.3 | 3.6 | 4.1 | 2.6 | 4.0 | 5.2 | 3.6 | 4.8 | NaN | 5.1 | 4.5 | 3.4 | 4.1 | 4.0 | 3.9 | 4.0 | 4.5 | 3.5 |
| IFJp – L | 3.6 | 3.3 | 4.2 | 4.1 | 4.3 | 5.2 | 3.7 | 4.0 | 4.3 | 4.2 | 4.4 | 4.9 | 3.8 | 3.8 | 4.1 | 4.0 | 3.7 | 4.2 | 4.9 | 4.2 | 5.0 | 5.4 | NaN | 4.6 | 4.0 | 4.4 | 4.4 | 4.1 | 4.3 | 4.8 | 4.1 |
| p9-46v | 4.3 | 3.5 | 4.3 | 3.0 | 4.4 | 4.5 | 2.6 | 3.8 | 3.6 | 4.2 | 4.1 | 4.5 | 3.0 | 3.6 | 3.4 | 3.1 | 3.5 | 4.5 | 4.1 | 3.5 | 4.0 | 4.8 | 4.6 | NaN | 2.8 | 4.0 | 3.5 | 4.4 | 3.2 | 3.9 | 3.1 |
| **46** | 4.6 | 4.1 | 3.4 | 3.3 | 2.7 | 3.7 | 2.5 | 4.2 | 4.0 | 4.5 | 3.9 | 3.3 | 2.5 | 4.4 | 3.3 | 2.6 | 4.4 | 3.8 | 2.9 | 3.7 | 2.5 | 3.4 | 4.0 | 2.6 | NaN | **3.4** | 4.2 | 3.8 | 4.3 | 3.0 | 4.2 |
| **LIPd –L** | 4.4 | 2.9 | 4.6 | 3.9 | 4.4 | 4.5 | 3.6 | 4.4 | 3.9 | 4.8 | 5.3 | 5.1 | 4.2 | 4.1 | 4.5 | 5.9 | 3.3 | 4.4 | 4.7 | 4.5 | 5.6 | 4.2 | 4.6 | 4.5 | **4.2** | NaN | 3.8 | 4.7 | 5.1 | 5.5 | 4.7 |

| | | | | | | | | | | | | | | | | | | | | | | | | | | | | | | | |
|---|---|---|---|---|---|---|---|---|---|---|---|---|---|---|---|---|---|---|---|---|---|---|---|---|---|---|---|---|---|---|---|
| FOP4 –L | 4.4 | 4.1 | 3.4 | 3.6 | 2.8 | 3.5 | 2.3 | 4.3 | 4.5 | 4.4 | 4.7 | 4.1 | 3.6 | 4.3 | 3.0 | 3.4 | 4.1 | 2.8 | 3.6 | 4.1 | 3.1 | 3.7 | 4.1 | 3.4 | 4.6 | 3.9 | NaN | 4.9 | 5.5 | 3.8 | 4.8 |
| AVI – L | 5.2 | 4.9 | 3.1 | 3.6 | 3.2 | 2.8 | 2.8 | 4.1 | 4.2 | 4.9 | 4.0 | 3.1 | 3.2 | 4.4 | 2.4 | 3.3 | 3.8 | 3.5 | 3.6 | 3.7 | 2.9 | 3.2 | 4.2 | 3.6 | 4.2 | 4.7 | 5.0 | NaN | 4.8 | 3.6 | 4.4 |
| PFt | 4.1 | 4.5 | 3.9 | 3.6 | 2.9 | 3.5 | 2.1 | 3.9 | 4.7 | 4.3 | 5.0 | 4.0 | 3.2 | 4.5 | 2.5 | 3.5 | 4.5 | 3.6 | 3.4 | 4.3 | 1.4 | 3.8 | 4.1 | 1.4 | 0.6 | 4.5 | 4.8 | 4.5 | NaN | 5.0 | 4.8 |
| IP2 – L | 4.5 | 2.9 | 5.0 | 4.0 | 4.1 | 5.3 | 3.5 | 4.4 | 3.0 | 4.3 | 4.5 | 5.8 | 4.0 | 3.6 | 4.1 | 6.1 | 2.7 | 4.5 | 4.7 | 3.8 | 5.3 | 4.9 | 4.8 | 4.5 | 3.3 | 4.9 | 3.4 | 4.0 | 5.0 | NaN | 3.1 |
| FOP5 | 4.4 | 4.8 | 3.3 | 4.3 | 3.4 | 2.7 | 2.2 | 4.4 | 4.4 | 4.3 | 4.4 | 3.4 | 3.1 | 3.5 | 2.5 | 3.4 | 4.5 | 2.5 | 4.2 | 4.8 | 3.2 | 4.0 | 4.4 | 3.5 | 4.4 | 4.5 | 5.4 | 5.0 | 4.5 | 3.4 | NaN |

Supplementary Table S4: Effective connectivity (DCM) for schizophrenia patients (N=66) during WM manipulation: T-values of effective connectivity across the 31 cortical parcels most engaged during WM manipulation. Rows represent connectivity "to" that region, while columns represent connectivity "from" that region. Regions are annotated according to the Human Connectome Project (HCP) parcellations(1). In bold are the left 46 and LIPd regions-of.-interest corresponding to previous connectivity findings associated with dysfunctional context updating and delusions(2) that are examined in the independent samples herein.

| | SCEF | a24pr | 44 – R | IFJp –R | IFSp | 46v | a10p | LIPd –R | FOP4 -R | AVI –R | AIP | IP2 – R | P10p | a32pr | SFL | 7PC | P32pr | 8BM | 44 – L | 47l | 6r | IFJa | IFJp –L | p9-46v | 46 | LIPd –L | FOP4 –L | AVI –L | PFt | IP2 – L | FOP5 |
|---|---|---|---|---|---|---|---|---|---|---|---|---|---|---|---|---|---|---|---|---|---|---|---|---|---|---|---|---|---|---|---|
| SCEF | NaN | 5.1 | 5.9 | 1.2 | 3.8 | 3.5 | 1.9 | 1.7 | 6.0 | 5.3 | 2.5 | 4.7 | 2.0 | 4.8 | 5.6 | 3.9 | 5.4 | 4.4 | 0.3 | 3.9 | 4.3 | 5.3 | 3.1 | 0.7 | 3.4 | 5.5 | 5.5 | 5.3 | 2.0 | 1.6 | 5.9 |
| a24pr | 5.7 | NaN | 6.1 | 5.1 | 4.3 | 4.9 | 1.8 | 5.3 | 5.4 | 5.1 | 5.6 | 5.0 | 3.6 | 5.6 | 4.3 | 4.1 | 4.9 | 4.7 | 5.1 | 3.8 | 5.8 | 5.5 | 4.6 | 3.4 | 3.3 | 5.1 | 5.2 | 4.9 | 5.8 | 5.2 | 5.5 |
| 44 –R | 4.5 | 5.3 | NaN | 3.8 | 2.8 | -0.2 | 1.8 | 5.1 | 5.4 | 1.5 | 1.6 | 0.3 | 1.7 | 5.2 | 1.1 | 3.6 | 5.9 | 4.3 | 0.8 | 3.8 | 0.6 | 3.1 | 0.9 | 3.7 | 4.4 | 5.6 | 1.0 | 5.4 | 5.0 | 4.9 | 1.3 |
| IFJp – R | 2.2 | 5.0 | 0.2 | NaN | 4.4 | 0.7 | 1.6 | 1.0 | 4.6 | 3.9 | 1.5 | 0.8 | 1.9 | 3.4 | 4.6 | 3.3 | 4.3 | 3.8 | 5.3 | 3.1 | 4.6 | 4.8 | 4.8 | 2.3 | 2.9 | 5.1 | 4.2 | 4.7 | 4.3 | 0.7 | 1.0 |
| IFSp | -0.2 | 3.8 | -0.4 | 0.4 | NaN | -0.2 | 1.1 | -1.0 | 0.8 | -0.2 | 1.3 | -0.7 | 1.3 | 3.5 | -1.0 | 0.2 | 0.4 | -1.2 | -1.0 | -0.1 | -1.6 | -1.0 | -1.4 | -0.2 | 3.0 | -0.4 | -0.4 | -1.2 | 0.6 | -0.9 | -0.2 |
| 46v | -0.5 | 4.3 | 0.1 | -0.5 | 4.0 | NaN | 1.9 | 0.1 | 4.8 | 4.9 | 0.5 | -0.2 | 1.1 | 3.6 | -0.6 | 2.8 | 4.7 | 3.2 | -0.4 | 3.5 | -0.8 | -0.6 | -0.2 | 0.3 | 1.1 | -0.1 | 2.1 | 0.4 | 5.6 | -1.0 | 0.8 |
| a10p | 0.7 | 0.9 | 0.7 | 0.7 | 0.2 | 0.6 | NaN | 1.2 | 1.6 | 0.4 | 1.2 | 0.7 | 1.5 | 1.3 | 1.1 | 1.3 | 0.0 | 0.1 | 1.2 | 2.2 | 1.6 | 0.5 | 1.1 | 1.2 | 1.4 | 0.8 | 0.2 | 0.2 | 2.3 | 1.5 | 1.0 |
| LIPd –R | 1.4 | 4.9 | 2.1 | 1.2 | 0.7 | 1.4 | 1.6 | NaN | 4.9 | 5.4 | 1.1 | 4.5 | 2.0 | 4.1 | 1.2 | 1.3 | 4.7 | 4.0 | 1.4 | 3.9 | 1.6 | 1.3 | 4.7 | 0.7 | 3.8 | 6.2 | 3.6 | 5.3 | 0.6 | 1.4 | 5.0 |
| FOP4 -R | 5.8 | 4.8 | 5.4 | 4.4 | 4.4 | 5.0 | 2.1 | 4.5 | NaN | 6.1 | 5.7 | 4.5 | 2.8 | 4.8 | 4.4 | 4.4 | 5.6 | 4.2 | 6.2 | 4.2 | 5.9 | 5.5 | 4.2 | 3.9 | 3.8 | 3.7 | 5.5 | 5.5 | 5.8 | 3.9 | 5.9 |
| AVI – R | 6.1 | 4.8 | 6.5 | 4.2 | 4.5 | 5.6 | 1.8 | 5.3 | 6.0 | NaN | 6.6 | 5.2 | 2.5 | 4.8 | 5.3 | 4.7 | 5.6 | 4.6 | 5.0 | 4.3 | 6.5 | 1.3 | 5.8 | 4.7 | 4.6 | 5.4 | 5.4 | 4.8 | 6.0 | 5.1 | 5.4 |
| AIP | 6.1 | 5.1 | 6.5 | 4.9 | 4.6 | 5.3 | 2.1 | 4.6 | 5.8 | 6.3 | NaN | 5.1 | 2.1 | 4.0 | 1.7 | 4.9 | 5.6 | 4.9 | 6.3 | 4.3 | 6.5 | 6.6 | 6.6 | 1.3 | 4.4 | 6.5 | 5.4 | 6.5 | 1.5 | 1.7 | 5.8 |
| IP2 – R | 4.8 | 4.8 | 4.9 | 4.1 | 3.9 | 4.3 | 1.9 | 1.5 | 4.8 | 5.1 | 1.8 | NaN | 2.6 | 3.8 | 4.9 | 3.8 | 4.6 | 4.0 | 4.3 | 4.0 | 5.9 | 5.2 | 5.1 | 4.2 | -0.3 | 5.4 | 5.5 | 4.9 | 4.8 | 0.2 | 2.4 |
| P10p | 2.0 | 2.9 | 1.8 | 1.5 | 1.8 | 1.5 | 2.3 | 1.3 | 1.9 | 1.8 | 1.9 | 2.1 | NaN | 3.0 | 1.5 | 1.4 | 2.6 | 1.3 | 1.7 | 1.6 | 1.6 | 1.2 | 1.8 | 0.9 | 1.4 | 1.3 | 1.7 | 0.8 | 2.5 | 1.4 | 0.7 |
| a32pr | 4.8 | 5.3 | 5.0 | 3.7 | 3.6 | 3.9 | 1.7 | 3.7 | 4.6 | 4.9 | 4.2 | 4.0 | 3.5 | NaN | 4.2 | 3.1 | 4.8 | 4.3 | 4.8 | 4.0 | 4.3 | 4.5 | 4.5 | 2.7 | 3.3 | 5.1 | 4.6 | 4.6 | 4.8 | 3.9 | 4.7 |
| SFL | 1.5 | 0.9 | 2.0 | 4.3 | 0.2 | 0.4 | 2.2 | 1.3 | 4.7 | 5.0 | 5.3 | 4.7 | 2.3 | 0.6 | NaN | 3.5 | 1.1 | -0.1 | 1.0 | 3.7 | 0.5 | 1.4 | 1.3 | 4.2 | 4.4 | 1.4 | 0.8 | 1.1 | 1.3 | 1.4 | 1.5 |
| 7PC | 4.8 | 3.8 | 1.2 | 0.5 | 0.7 | 0.3 | 2.5 | 1.2 | 4.4 | 4.3 | 1.7 | 3.5 | 2.3 | 3.4 | 4.0 | NaN | 4.4 | 0.7 | 0.4 | 2.9 | 1.3 | 0.8 | 0.9 | 0.3 | 3.8 | 0.9 | 2.8 | 4.4 | 5.2 | 1.0 | 3.5 |
| P32pr | 5.6 | 5.3 | 6.5 | 4.5 | 3.8 | 5.6 | 1.7 | 4.9 | 5.5 | 6.1 | 6.0 | 4.6 | 3.0 | 4.9 | 5.3 | 4.9 | NaN | 5.1 | 6.8 | 4.8 | 6.8 | 5.9 | 6.3 | 4.4 | 4.1 | 6.1 | 6.2 | 6.4 | 6.3 | 5.5 | 6.1 |
| 8BM | 1.2 | 0.9 | 1.9 | 0.8 | -0.5 | 4.2 | 1.1 | 1.1 | 4.2 | 5.1 | 4.9 | 4.2 | 1.8 | 1.1 | 1.2 | 4.0 | 1.0 | NaN | 0.5 | 3.6 | 1.5 | -0.4 | 1.7 | 3.6 | 3.0 | 1.2 | 0.6 | 1.1 | 1.1 | 1.0 | 1.2 |
| 44 –L | 0.3 | 1.1 | 0.5 | 1.6 | 0.9 | 0.5 | 2.3 | 1.6 | 5.7 | 1.0 | 2.3 | -0.7 | 2.0 | 1.1 | 0.6 | 3.3 | 6.2 | -0.2 | NaN | 4.5 | -0.9 | -0.4 | -0.6 | 1.2 | 4.8 | 1.3 | 4.4 | 1.3 | 1.8 | 0.2 | 1.6 |
| 47l | 0.6 | 0.5 | 1.3 | 3.4 | 0.2 | 3.8 | 2.2 | 3.2 | 4.1 | 4.4 | 4.0 | 3.4 | 1.7 | 0.6 | 0.7 | 3.1 | 4.5 | 0.9 | 4.6 | NaN | 1.4 | 0.8 | 1.2 | 3.9 | 3.5 | 0.6 | 4.0 | 0.0 | 0.7 | 0.7 | 4.6 |
| 6r | 0.0 | 5.1 | 0.1 | 0.3 | -0.1 | 0.9 | 2.2 | 0.0 | 5.8 | 6.0 | 1.4 | 2.3 | 0.0 | 5.1 | -0.3 | 0.4 | 5.8 | -0.5 | -0.7 | 1.4 | NaN | -0.2 | -0.4 | -0.1 | 1.4 | 5.5 | 1.8 | 1.3 | 2.1 | 1.4 | 0.7 |
| IFJa | 1.6 | 5.4 | 0.6 | 1.7 | 0.9 | 1.0 | 1.6 | 1.6 | 5.8 | 4.9 | 2.6 | 1.7 | -0.1 | 4.8 | 0.6 | 4.1 | 5.6 | 2.5 | 4.7 | 4.3 | 5.2 | NaN | 1.9 | 2.4 | 1.0 | 5.9 | 3.9 | 5.0 | 5.7 | 1.4 | 1.6 |
| IFJp – L | 0.5 | 1.1 | 1.6 | 0.6 | 0.9 | 1.3 | 1.9 | 1.2 | 4.7 | 2.7 | 3.1 | 2.4 | 0.5 | 4.5 | 0.6 | 0.9 | 6.0 | 0.4 | 2.2 | 1.5 | 1.2 | 0.8 | NaN | 1.3 | 1.2 | 1.8 | 1.8 | 1.6 | 5.6 | 0.8 | 6.4 |
| p9-46v | 0.2 | 3.0 | 0.2 | -0.8 | 2.3 | 0.1 | 2.0 | 0.3 | 3.6 | 4.7 | 0.8 | 1.0 | 1.0 | 2.6 | -0.3 | 3.7 | 3.6 | 3.1 | -0.6 | 3.3 | -0.3 | -0.7 | 0.6 | NaN | -0.4 | 2.9 | 4.3 | 0.4 | 3.1 | 0.4 | -0.2 |

| | | | | | | | | | | | | | | | | | | | | | | | | | | | | | | | |
|---|---|---|---|---|---|---|---|---|---|---|---|---|---|---|---|---|---|---|---|---|---|---|---|---|---|---|---|---|---|---|---|
| 46 | 3.4 | 3.5 | 4.3 | 2.6 | 3.1 | -0.1 | 1.9 | 4.1 | 3.7 | 4.1 | 4.6 | 1.9 | 1.7 | 3.0 | 3.3 | 3.7 | 3.7 | 2.7 | 0.8 | 3.5 | -0.1 | 3.0 | 2.3 | 3.8 | NaN | 4.0 | 3.7 | 4.4 | 3.9 | 3.9 | 4.1 |
| LIPd –L | 2.5 | 4.9 | 2.7 | 2.0 | 1.1 | 1.8 | 1.9 | 2.5 | 5.2 | 5.9 | 6.9 | 5.3 | 1.6 | 4.8 | 1.9 | 4.0 | 5.6 | 1.4 | 1.9 | 4.0 | 2.9 | 2.4 | 2.4 | 1.5 | 3.9 | NaN | 4.9 | 6.1 | 2.5 | 5.8 | 5.8 |
| FOP4 –L | 5.6 | 4.6 | 4.6 | 4.1 | 3.0 | 5.3 | 1.4 | 4.8 | 5.3 | 5.8 | 5.6 | 5.4 | 1.4 | 4.4 | 4.1 | 3.7 | 5.6 | 0.8 | 1.9 | 3.9 | 2.1 | 1.2 | 5.7 | 4.6 | 4.0 | 5.5 | NaN | 5.5 | 4.7 | 5.0 | 5.8 |
| AVI – L | 1.4 | 4.4 | 6.0 | 4.8 | 3.1 | 4.4 | 1.4 | 5.3 | 5.1 | 5.8 | 6.1 | 5.1 | 1.5 | 4.3 | 3.7 | 3.5 | 5.8 | 3.6 | 5.4 | 4.6 | 5.6 | 5.2 | 1.8 | 4.7 | 4.2 | 6.0 | 5.7 | NaN | 6.0 | 1.1 | 5.8 |
| PFt | 6.7 | 4.9 | 2.2 | 4.2 | 0.9 | 1.8 | 3.2 | 5.5 | 5.6 | 5.9 | 6.1 | 5.6 | 2.5 | 4.7 | 5.2 | 3.9 | 5.5 | 4.8 | 5.2 | 4.2 | 6.6 | 5.7 | 6.1 | 3.3 | 4.7 | 5.8 | 4.2 | 6.0 | NaN | 5.5 | 5.8 |
| IP2 – L | 1.9 | 5.2 | 2.4 | 1.7 | 1.1 | 1.0 | 2.0 | 2.0 | 4.9 | 5.3 | 5.0 | 4.1 | 2.4 | 4.0 | 1.6 | 1.8 | 4.8 | 1.1 | 1.4 | 3.8 | 2.2 | 1.8 | 1.7 | 1.1 | 3.4 | 1.9 | 5.0 | 5.2 | 2.2 | NaN | 5.4 |
| FOP5 | 5.7 | 5.2 | 5.9 | 3.9 | 4.1 | 4.8 | 2.0 | 1.6 | 5.4 | 5.7 | 2.4 | 1.7 | 2.3 | 5.0 | 4.8 | 3.5 | 6.2 | 4.4 | 5.5 | 4.8 | 6.2 | 5.8 | 5.9 | 4.6 | 4.5 | 5.8 | 5.7 | 5.4 | 5.6 | 4.8 | NaN |

Supplementary Table S5: Effective connectivity (DCM) for schizophrenia patients (N=66) during WM maintenance: T-values of effective connectivity across the 31 cortical parcels most engaged during WM maintenance. Rows represent connectivity "to" that region, while columns represent connectivity "from" that region. Regions are annotated according to the Human Connectome Project (HCP) parcellations(1).

Supplementary References:


1. Glasser MF*, et al.* (2016) A multi-modal parcellation of human cerebral cortex. *Nature* 536:171.
2. Kaplan C.M.*, et al.* (2016) Estimating changing contexts in schizophrenia. *Brain* 139:2082-2095.
